# The minus end depolymerase KIF2A drives flux-like treadmilling of γTuRC-uncapped microtubules

**DOI:** 10.1101/2023.04.06.535808

**Authors:** Gil Henkin, Cláudia Brito, Claire Thomas, Thomas Surrey

## Abstract

During mitosis, a functional spindle requires high microtubule turnover. Such turnover is highlighted by the multiple functions of spindle poles, where microtubule minus ends are concentrated, and where microtubule nucleation and depolymerization happen side by side. How these seemingly antagonistic processes are coordinated during poleward microtubule flux is not understood. Here we reconstitute this coordination *in vitro* combining different pole localized activities. We find that the spindle pole-localized kinesin-13 KIF2A is a microtubule minus-end depolymerase, in contrast to its paralog MCAK. Due to its asymmetric activity, KIF2A still allows microtubule nucleation by plus-end growth from the γ-tubulin ring complex (γTuRC), which in turn serves as a protective cap that shields the minus end against KIF2A binding. Efficient γTuRC-uncapping requires the combined action of KIF2A and a microtubule severing enzyme, then leading to treadmilling of the uncapped microtubule driven by KIF2A. Together these results provide insight into the molecular mechanisms by which a minimal protein module coordinates microtubule nucleation and depolymerization at spindle poles consistent with their role in poleward microtubule flux.

## INTRODUCTION

The microtubule cytoskeleton is essential for a multitude of processes in eukaryotic cells, such as intracellular organization and trafficking, cell division and differentiation. Microtubules are structurally polar filaments with two biochemically distinct ends with characteristic differences in their dynamic properties that are critical for microtubule function (Akhmanova and Steinmetz, 2015; Gudimchuk and McIntosh, 2021). Both ends of microtubules can switch stochastically between episodes of growth and shrinkage, a property called dynamic instability, which is ultimately a consequence of GTP hydrolysis at the microtubule end (Desai and Mitchison, 1997; Mitchison and Kirschner, 1984). The dynamic properties of microtubule plus ends and their regulation have been extensively studied, both in cells and also *in vitro* with purified proteins, but much less is known about the regulation of microtubule minus end dynamics (Akhmanova and Steinmetz, 2019).

Across eukaryotic cells, major microtubule nucleation pathways require the γ-tubulin ring complex (γTuRC), a large multi-subunit protein complex that serves as a template for microtubule formation (Kollman et al., 2011; Tovey and Conduit, 2018). Nucleation from purified γTuRC *in vitro* results in microtubules with stably γTuRC-capped minus ends and dynamic plus ends (Berman et al., 2023; Consolati et al., 2020; Rai et al., 2022; Thawani et al., 2020; Wieczorek et al., 2021). To what extent γTuRC-nucleated microtubule minus ends remain capped in cells is not clear, and recent electron microscopy data suggests there is a mixture of capped and uncapped minus ends in cells (Kiewisz et al., 2022; Laguillo-Diego et al., 2023).

In mitotic and meiotic spindles during cell division, microtubule minus ends at the spindle poles appear to slowly depolymerize while dynamic microtubule plus ends in the spindle center display net polymerization, leading to a constant flux of microtubule mass from the spindle center to its poles (Barisic et al., 2021; Cameron et al., 2006; Ferenz and Wadsworth, 2007; Ganem et al., 2005; Mitchison, 1989; Risteski et al., 2022; Rogers et al., 2005; Steblyanko et al., 2020). How microtubule minus end dynamics at spindle poles are regulated during poleward flux remains a major open question. On one hand, γTuRC is enriched at spindle poles, in agreement with the poles’ microtubule-nucleating properties. On the other hand, a number of microtubule-destabilizing proteins involved in regulating microtubule flux are also known to accumulate at spindle poles, such as the microtubule depolymerase KIF2A, a kinesin-13 (Gaetz and Kapoor, 2004; Ganem and Compton, 2004; Welburn and Cheeseman, 2012), and the microtubule severases like katanin, fidgetin and spastin (Jiang et al., 2017; McNally et al., 1996; Zhang et al., 2007). The molecular mechanism by which this mixture of seemingly antagonistic microtubule nucleating and destabilizing activities at the spindle pole can control microtubule minus end dynamics to ensure correct microtubule flux and hence correct spindle function is unknown.

The kinesin-13s are not directionally motile, but instead bind microtubules in a diffusive manner and depolymerize them from their ends. This function requires ATPase activity as well as the positively-charged and class-specific “neck”, which is directly N-terminal to the conserved kinesin motor domain (Desai et al., 1999; Ems-McClung and Walczak, 2010; Friel and Welburn, 2018; Moores and Milligan, 2006). The N and C termini of the kinesin-13s drive cellular localization and homodimerization, respectively (Maney et al., 2001; Welburn and Cheeseman, 2012). In the ATP state both the neck and motor domain bind two longitudinally bound tubulin dimers in a bent conformation, which suggests an ‘unpeeling’ mechanism to remove tubulin from microtubule ends (Benoit et al., 2018; Trofimova et al., 2018; Wang et al., 2017).

The majority of kinesin-13 studies have focused on MCAK (KIF2C), which preferentially localizes to kinetochores in cells and is involved in correct chromosome-spindle attachment (Ganem et al., 2005; Manning et al., 2007; Welburn and Cheeseman, 2012; Wordeman et al., 2007). *In vitro*, purified MCAK slowly depolymerizes both ends of artificially stabilized microtubules at similar speeds, and promotes the transition from growth to fast shrinkage, called catastrophe, of dynamic microtubules (Desai et al., 1999; Gardner et al., 2011; Montenegro Gouveia et al., 2010). The paralog KIF2A on the other hand, localizes mainly at spindle poles, and is important for correct spindle architecture and length, promoting poleward flux (Gaetz and Kapoor, 2004; Ganem and Compton, 2004; Steblyanko et al., 2020; Uehara et al., 2013; Wilbur and Heald, 2013). Although KIF2A has been studied less *in vitro* (Desai et al., 1999; Wilbur and Heald, 2013), it is thought that KIF2A and MCAK share identical enzymatic activities, with their functional differences in cells resulting from different localizations mediated by specific binding partners. It is however still unknown how KIF2A contributes to the control of microtubule minus end dynamics at spindle poles, especially given that most minus ends at spindle poles are expected to be capped after having been nucleated by γTuRCs.

Microtubule uncapping by microtubule severases could allow KIF2A to depolymerize minus ends at spindle poles (Sharp and Ross, 2012; Zhang et al., 2007), and indeed severases promote poleward flux (Guerreiro et al., 2021; Huang et al., 2021). They are hexameric AAA proteins that pull on the negatively charged C-terminal tails of tubulin on the microtubule lattice, inducing damage that can lead to microtubule severing, exposing new plus and minus ends (Roll-Mecak and Vale, 2008; Zehr et al., 2020). This process has however been shown to be inefficient in the presence of soluble tubulin, due to spontaneous re-incorporation of tubulin at the damage sites (Jiang et al., 2017; Vemu et al., 2018), raising the question of how efficient minus end uncapping by severases could be at the spindle poles. The molecular mechanisms of how γTuRC, KIF2A and severases in combination control microtubule minus end dynamics to ensure correct spindle function remain largely unknown.

Here, we demonstrate that KIF2A is an inherently asymmetric microtubule depolymerase. It preferentially binds to and depolymerizes the minus ends of microtubules, in agreement with its function in the spindle, and distinct from the symmetric behavior of its paralog MCAK. KIF2A alone is sufficient to induce microtubule treadmilling that recapitulates poleward flux, by slowly depolymerizing the minus ends of dynamic microtubules while permitting plus end growth. We find that KIF2A still allows microtubule nucleation by γTuRC, and that γTuRC capping of minus ends interferes with KIF2A-mediated minus end depolymerization. The severase spastin and KIF2A are together required to create fresh minus ends via active γTuRC-uncapping and to induce KIF2A-mediated microtubule treadmilling. Altogether, our results provide new insight into the molecular mechanism of the control of microtubule minus end dynamics by the γTuRC/KIF2A/severase module, consistent with their role at spindle poles during cell division.

## RESULTS

### KIF2A is a microtubule minus end depolymerase

We expressed and purified the human kinesin-13 KIF2A fused to the fluorescent protein mScarlet (mScarlet-KIF2A; Fig. 1A, Suppl. Fig. 1A) and studied its depolymerization activity on microtubules using a total internal reflection fluorescence (TIRF) microscopy-based *in vitro* assay (Fig. 1A). We were surprised to observe strongly asymmetric depolymerization of glass-immobilized microtubules that were stabilized using the slowly hydrolyzable GTP analog GMPCPP (Movie 1). One microtubule end depolymerized considerably faster than the other (Fig. 1B), and mScarlet-KIF2A strongly accumulated at the more-quickly shrinking end (Suppl. Fig. 2A). This is in contrast to the activity of other kinesin-13s such as MCAK, which display rather symmetric depolymerization activity at both ends of stabilized microtubules (Desai et al., 1999; Helenius et al., 2006; Hunter et al., 2003).

**Fig. 1.**
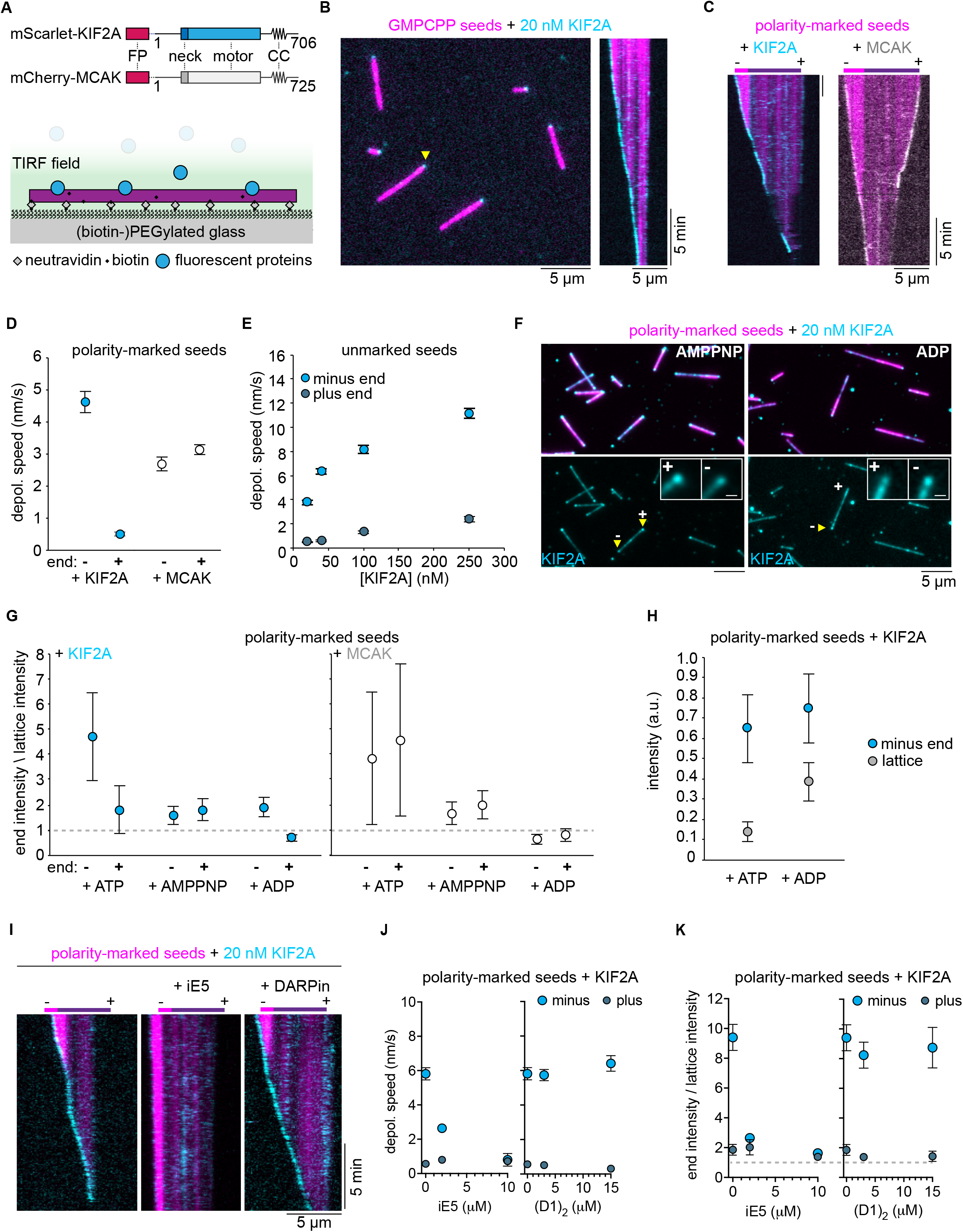
The depolymerase KIF2A preferentially accumulates and depolymerizes the minus ends of microtubules. **A** Schematic of fluorescently-tagged KIF2A and MCAK constructs used in this study, indicating the fluorescent protein (“FP”) fused via a flexible linker to the N terminus of each protein, the conserved internal motor domain, and the class-specific “neck” and predicted coiled-coil dimerization domain (“CC”) N and C terminal to the motor domain. Below, schematic of TIRF microscopy experiment set-up. **B** TIRF microscopy image of 20 nM mScarlet-KIF2A (cyan) binding to GMPCPP-microtubule seeds (AlexaFluor647, 5%; magenta) bound to the glass surface. A kymograph of a depolymerizing microtubule seed indicated by the yellow arrow is on the right. **C** Kymographs of GMPCPP-microtubule seeds with a dimly labeled plus end and brightly labeled minus end (AlexaFluor647; magenta) being depolymerized by 20 nM mScarlet-KIF2A (left; cyan) or 20 nM mCherry-MCAK (right; gray). **D** Depolymerization rates of each end of polarity-marked GMPCPP seeds by 20 nM mScarlet-KIF2A (*n* = 29 microtubules) and 20 nM mCherry-MCAK (*n* = 33 microtubules). Error bars are SEM. **E** Depolymerization rates of GMPCPP-microtubule seeds at varying concentrations of mScarlet-KIF2A. Polarity of the seed was assigned assuming the minus end depolymerizes at the faster rate. Number of microtubule seeds measured per condition: mScarlet-KIF2A: 20 nM, *n* = 46; 40 nM, *n* = 61; 100 nM, *n* = 78; 250 nM, *n* = 64. Error bars are SEM. **F** Localization of mScarlet-KIF2A (20 nM) on polarity-marked microtubule seeds in AMPPNP and ADP. Images are averages of 10 frames. Top row: merged channels (AlexaFluor647 tubulin, magenta; mScarlet-KIF2A, cyan). Bottom row, mScarlet-KIF2A channel alone. Bottom inset, cropped and enlarged images of mScarlet-KIF2A binding to microtubule ends, showing its localization to both ends in AMPPNP, and its localization to the minus end, and not the plus end, in ADP. Inset scale bars: 1 µm. **G** Ratio of intensity of mScarlet-KIF2A or mCherry-MCAK (both 20 nM) at microtubule ends to their intensity on the microtubule lattice (see Methods). Dashed line at a ratio of 1 indicates equal binding to the end and the lattice. Number of microtubule seeds measured per condition: mScarlet-KIF2A: ATP, *n* = 26; AMPPNP, *n* = 29; ADP, *n* = 36: mCherry-MCAK: ATP, *n* = 47; AMPPNP, *n* = 40; ADP, *n* = 32. Error bars are SEM. **H** “Raw” mScarlet-KIF2A intensities at minus ends and on the lattice in ATP and ADP (see Methods). Error bars are SEM. **I** Kymographs of polarity-marked GMPCPP seeds (Atto647; magenta) in the presence of 20 nM mScarlet-KIF2A (cyan) and either 2 µM iE5 (middle) or 15 µM (D1)_2_ (right). **J** Depolymerization speeds of plus and minus ends of polarity-marked GMPCPP seeds in the presence of 20 nM mScarlet-KIF2A (*n* = 173 microtubules) and either iE5 (2 µM, *n* = 149 microtubules; 10 µM, n = 125) or (D1)_2_ (3 µM, *n* = 149 microtubules; 15 µM, *n* = 141 microtubules). Error bars are SEM. **K** Ratio of microtubule end and lattice intensities of mScarlet-KIF2A (*n* = 44 microtubules) in the absence or presence of iE5 (2 µM, *n* = 44 microtubules; 10 µM, n = 55) or (D1)_2_ (3 µM, *n* = 52 microtubules; 15 µM, *n* = 37 microtubules) at the indicated concentrations. Dashed line indicates equal binding to the end and the lattice (ratio = 1). Error bars are SEM.

Polarity-labeling of GMPCPP-microtubules showed that mScarlet-KIF2A preferentially bound to and depolymerized microtubule minus ends; plus ends depolymerized between 5 and 10 times more slowly (Fig. 1C left, Fig. 1D). A purified KIF2A construct without a fluorescent tag (Suppl. Fig. 1B) showed similar asymmetric depolymerase behavior on polarity-marked GMPCPP-microtubules (Suppl. Fig. 2B, C). To be able to directly compare the activities of KIF2A and MCAK under the same assay conditions, we expressed and purified MCAK fused to mCherry (mCherry-MCAK; Fig. 1A, Suppl. Fig. 1C) and confirmed that MCAK does not have a minus end preference, neither for accumulation nor for depolymerization of GMPCPP-microtubules (Fig. 1C right, Fig. 1D).

Varying the mScarlet-KIF2A concentration, we observed that the depolymerization speed of both the minus and the plus end of non-polarity-marked microtubules (distinguished by considering that the minus end depolymerizes faster) increased with the KIF2A concentration, maintaining a strongly asymmetric depolymerization character at all the KIF2A concentrations tested (Fig. 1E). All-in-all, this demonstrates that KIF2A is inherently different from MCAK, preferentially depolymerizing minus ends, consistent with its proposed role at spindle poles (Cameron et al., 2006; Gaetz and Kapoor, 2004; Ganem and Compton, 2004).

Kinesin-13s may recognize a particular lattice curvature at microtubule ends induced by ATP binding of the motor domain (Benoit et al., 2018; Desai et al., 1999; Hunter et al., 2003; Trofimova et al., 2018; Wang et al., 2017). Therefore, we asked if also the preferential accumulation of KIF2A at minus ends was dependent on its nucleotide state. We found that mScarlet-KIF2A accumulated at both ends of microtubules in the presence of the non-hydrolysable ATP analog AMPPNP (Fig. 1F, left), as observed previously with kinesin-13s (Desai et al., 1999; McHugh et al., 2019; Moores et al., 2002). However, we were surprised to find that in the presence of ADP KIF2A accumulated only at the minus and not at the plus ends (Fig. 1F, right). We quantified the extent of end versus lattice accumulation by measuring fluorescence intensity ratios. In the presence of ATP, KIF2A showed a marked binding preference for the minus end, but also accumulated mildly at the plus end (Fig. 1G, left, ATP), consistent with the observed difference in depolymerization rates at the two ends. In the presence of AMPPNP, KIF2A accumulated equivalently mildly at both plus and minus ends, remarkably without a minus end preference (Fig. 1G, left, AMPPNP), whereas with ADP KIF2A accumulation only showed a minus end preference (Fig. 1G, left, ADP). Because ADP also increased KIF2A’s binding to the microtubule lattice (Fig. 1H), the ratio of end-to-lattice intensity was lower for ADP than ATP, but the amount of KIF2A motors bound to minus ends was similar in the presence of ATP and ADP (Fig. 1H). Taken together, these results indicate that the selective microtubule minus end accumulation by KIF2A is distinct from the indiscriminate end accumulation of MCAK and that there is a structure at minus ends, and not at plus ends, that KIF2A can recognize independent of ATP binding. In contrast, MCAK did not show any end accumulation in the presence of ADP, whereas it accumulated at both ends equally strongly in ATP and equally mildly in AMPPNP (Fig. 1G, right), as observed previously (Desai et al., 1999; Helenius et al., 2006; Hertzer et al., 2006; McHugh et al., 2019). To further investigate the mechanism of minus end recognition by KIF2A, we next purified the microtubule end capping protein iE5 (Suppl. Fig. 1D) and DARPin (D1)_2_ (Suppl. Fig. 1E) which specifically cap either the minus or the plus ends of microtubules, respectively (Campanacci et al., 2019; Pecqueur et al., 2012). Using polarity-marked microtubules we observed that while (D1)_2_ had no influence on KIF2A’s capacity to accumulate at and depolymerize microtubule minus ends, as one would expect, iE5 effectively hindered both minus end accumulation and depolymerization by KIF2A (Fig. 1I, J and K). These results suggest that KIF2A might interact with the exposed surface of the α-tubulins that iE5 binds to, suggesting a new model for microtubule minus end recognition by KIF2A.

### KIF2A selectively depolymerizes the minus ends of dynamic microtubules

Having established a robust minus-end preference of KIF2A using stabilized microtubules, we next investigated if KIF2A also acts asymmetrically on dynamic microtubules growing from immobilized GMPCPP microtubule seeds in the presence of soluble GTP-tubulin (Fig. 2A; Movie 2). Dynamic microtubule extensions could be distinguished from seeds by including a higher ratio of fluorescent tubulin in the seed (Fig. 2B). In the absence of mScarlet-KIF2A, the faster growing microtubule plus ends could easily be distinguished from the more slowly growing minus ends (Fig. 2B – 0 nM KIF2A). Adding mScarlet-KIF2A selectively suppressed minus end growth already at low concentrations (Fig. 2B – 2 nM, 5 nM KIF2A) and caused selective minus-end depolymerization of the GMPCPP seed at higher concentrations. At the same time, plus end growth could be observed up to 100 nM of KIF2A (Fig. 2B – 20 nM, 100 nM KIF2A) without affecting the plus end growth speed (Fig. 2C top left). The plus end growth speed was also unaffected by MCAK (Fig. 2C top right), in agreement with previous MCAK studies (Gardner et al., 2011; Montenegro Gouveia et al., 2010). At higher concentrations, KIF2A also accumulates along the microtubule lattice, triggering more frequently the transition from plus end growth to shrinkage, thus increasing the plus end catastrophe frequency (Fig. 2B, C bottom left).

**Fig. 2.**
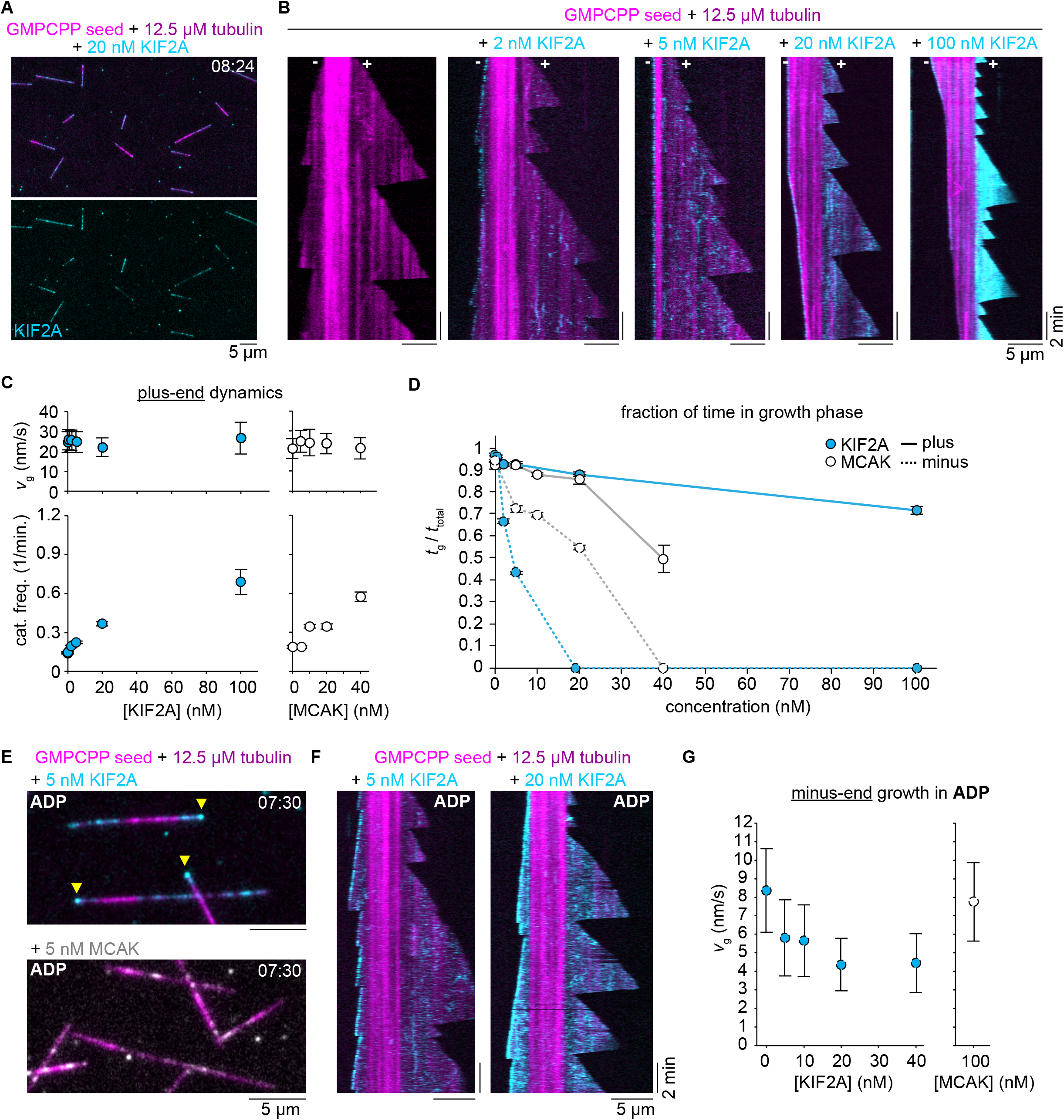
Preferential destabilization of dynamic microtubule minus ends by KIF2A. **A** Snapshot of dynamic microtubules polymerizing from brightly labeled GMPCPP seeds (Alexa647 tubulin, magenta) in the presence of mScarlet-KIF2A (cyan). Below, mScarlet-KIF2A channel alone. **B** Sequence of kymographs for representative microtubules polymerizing from bright seeds (Alexa647 tubulin, magenta) in the presence of increasing concentrations of mScarlet-KIF2A (cyan). mScarlet-KIF2A channels were additionally background subtracted and contrast was independently adjusted for each condition to show minus-end binding at a similar brightness. **C** Dynamics of microtubule plus-ends growing in the presence of increasing concentrations of mScarlet-KIF2A (left) or mCherry-MCAK (right), with a tubulin concentration of 12.5 µM. Both proteins show no major impact on growth speed (top) and a similar increase in catastrophe rate (bottom) with increasing concentrations. Number of growth speeds measured per condition: mScarlet-KIF2A: 0 nM, *n* = 502; 0.5 nM, *n* = 379; 2 nM, *n* = 508; 5 nM, *n* = 405; 20 nM, *n* = 572; 100 nM, *n* = 259: mCherry-MCAK: 0 nM, *n* = 275; 5 nM, *n* = 275; 10 nM, *n* = 406; 20 nM, *n* = 317; 40 nM, *n* = 164. Number of catastrophe rates measured per condition: mScarlet-KIF2A: 0 nM, *n* = 127; 0.5 nM, *n* = 90; 2 nM, *n* = 104; 5 nM, *n* = 71; 20 nM, *n* = 64; 100 nM, *n* = 26: mCherry-MCAK: 0 nM, *n* = 57; 5 nM, *n* = 60; 10 nM, *n* = 61; 20 nM, *n* = 41; 40 nM, *n* = 14. Error bars are SD for growth speed and SEM for catastrophe frequency. **D** Fraction of time in growth phase for minus and plus ends of dynamic microtubules in the presence of increasing concentrations of mScarlet-KIF2A or mCherry-MCAK and 12.5 µM tubulin. Lines are drawn as a guide to the eye. Number of minus end traces measured per condition: mScarlet-KIF2A: 0 nM, *n* = 127; 0.5 nM, *n* = 99; 2 nM, *n* = 60; 5 nM, *n* = 63: mCherry-MCAK: 0 nM, *n* = 59; 5 nM, *n* = 21; 10 nM, *n* = 33; 20 nM, *n* = 35. Number of plus end traces measured per condition: mScarlet-KIF2A: 0 nM, *n* = 131; 0.5 nM, *n* = 93; 2 nM, *n* = 95; 5 nM, *n* = 70; 20 nM, *n* = 67; 100 nM, *n* = 32: mCherry-MCAK: 0 nM, *n* = 58; 5 nM, *n* = 65; 10 nM, *n* = 67; 20 nM, *n* = 49; 40 nM, *n* = 34. Measurements of 0% were inferred by manual inspection of conditions in which seed ends only depolymerized. Error bars are SEM. **E** Snapshot of microtubules (magenta) growing from brightly-labeled seeds in the presence of mScarlet-KIF2A (top, cyan) or mCherry-MCAK (bottom, gray) in the presence of ADP and absence of ATP. Yellow arrowheads indicate mScarlet-KIF2A accumulated at the slower growing minus ends; mCherry-MCAK did not significantly decorate either end of the growing microtubule in the presence of ADP. Images are averages of two frames taken 5 seconds apart. **F** Representative kymographs showing minus end tracking of ADP-bound mScarlet-KIF2A (cyan) and partially suppressed minus end growth of microtubules growing from brightly-labeled seeds (magenta). **G** Microtubule minus-end growth speeds in the presence of increasing concentrations of mScarlet-KIF2A (left) or 100 nM mCherry-MCAK (right) in ADP. Number of growth speeds measured per condition: mScarlet-KIF2A: 0, *n* = 404; 5, *n* = 303; 10, *n* = 330; 20, *n* = 336; 40, *n* = 397: mCherry-MCAK: 100, *n* = 323. Error bars are SD.

The fraction of time spent growing is a convenient single parameter to capture differences in the dynamic state of microtubule plus and minus ends. For KIF2A it highlights a strong asymmetry between plus and minus end behavior across all KIF2A concentrations studied (Fig. 2D). In contrast, in the presence of MCAK we observed only a very mild asymmetry for the fraction of time plus and minus ends spend growing (Fig. 2D). These results establish that KIF2A also has a strong minus-end preference for dynamic microtubules and allows plus ends to grow up to quite high KIF2A concentration, which is characteristically distinct from its paralog MCAK.

Having observed that KIF2A preferentially accumulates at the minus ends of stabilized microtubules in the enzymatically inactive ADP-bound state (Fig. 1F, G), we wondered what effect ADP-bound KIF2A has on dynamic microtubule minus ends. We found that ADP-KIF2A remarkably accumulated at growing minus ends, but not at plus ends, in contrast to MCAK that did not visibly accumulate at any dynamic end (Fig. 2E, F). Minus end tracking by ADP-KIF2A slowed down, but did not completely block minus end growth (Fig. 2F, G), an effect not observed with MCAK (Fig. 2G). This shows that the mechanism by which KIF2A recognizes minus ends in the ADP state partially interferes with tubulin addition to the growing minus end, but that complete suppression of growth requires ATP-dependent enzymatic kinesin activity.

### KIF2A drives treadmilling of dynamic microtubules

Next, we wondered what would happen at increased tubulin concentrations, where microtubules grow faster and experience fewer catastrophes. This condition would allow KIF2A to depolymerize the entire stabilized seed and then to reach the minus end of the GDP lattice formed by previous plus end growth from the seed. Such a GDP-minus end may represent a close mimic of a microtubule end that KIF2A interacts with in a cell. We found that instead of inducing a stereotypical minus end catastrophe, the GDP-minus end underwent relatively slow depolymerization in the presence of either untagged KIF2A or mScarlet-KIF2A (Fig. 3A, B; Movie 3). Accumulated mScarlet-KIF2A visibly tracks the depolymerizing minus end, with occasional phases of more rapid shrinkage during which KIF2A was not accumulated (Fig. 3B; Movie 3). As the plus end continued to grow, this resulted in treadmilling of the microtubule across the glass surface, held to the surface by the crowding agent methylcellulose, until shrinkage from one or both ends completely depolymerized the microtubule or it became sufficiently small to diffuse away. This behavior is similar to a recent study reproducing microtubule treadmilling using MCAK and a combination of several other microtubule binding proteins that increased the plus end stability and growth speed to counteract MCAK’s destabilizing activity at plus ends (Arpağ et al., 2020). However, to our knowledge, our experiments with KIF2A are the first demonstration of treadmilling of a dynamic microtubule induced by a single microtubule binding protein. The treadmilling activity was robust over a range of KIF2A concentrations (Fig. 3B), and minus ends depolymerized in a KIF2A concentration dependent manner at rates from 10 to 30 nm/s, consistent with cellular rates of poleward flux (Cameron et al., 2006; Steblyanko et al., 2020), and more than an order of magnitude slower than depolymerization episodes in the absence of KIF2A (Fig. 3C).

**Fig. 3.**
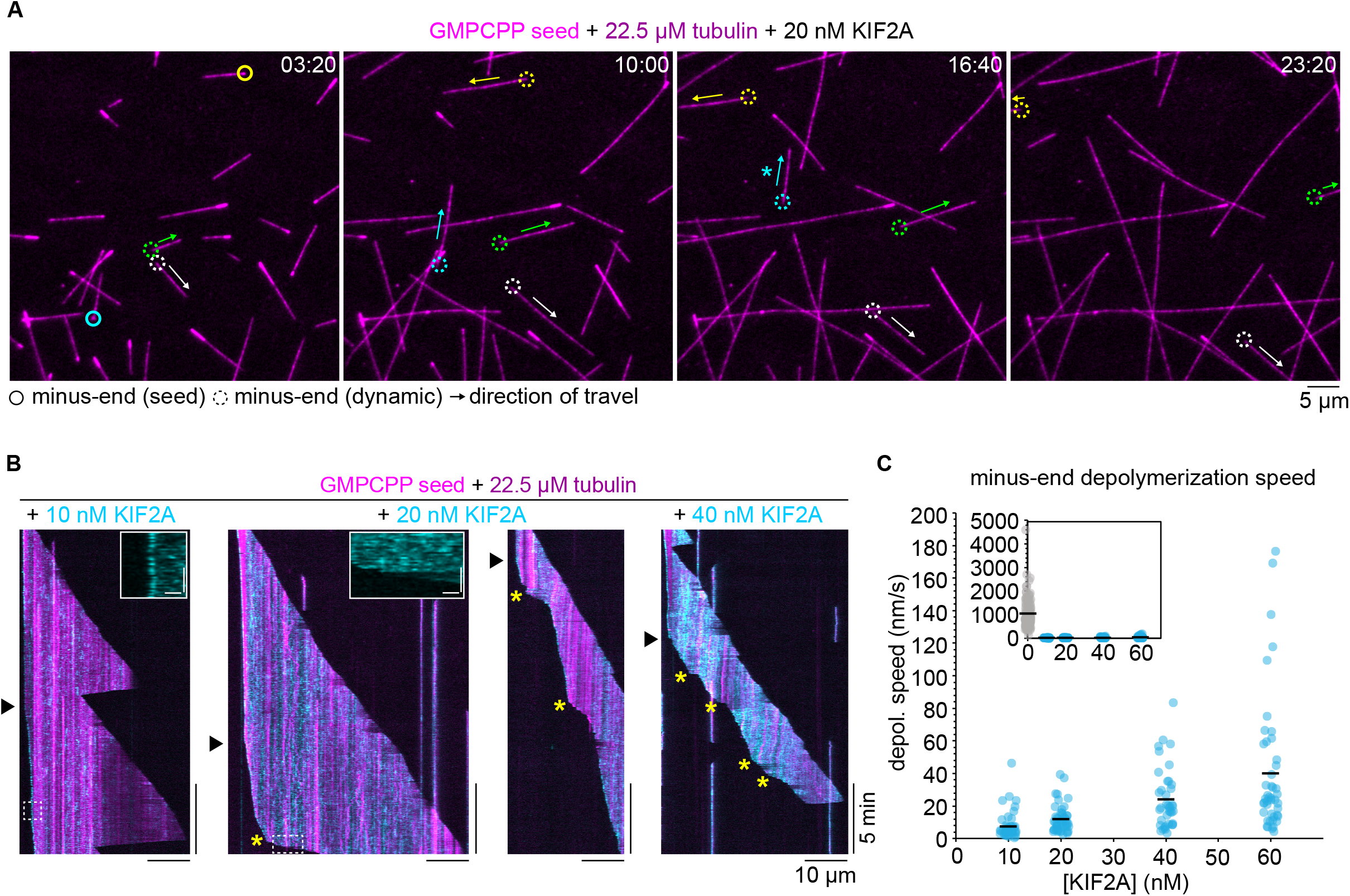
Treadmilling of dynamic microtubules driven by KIF2A. **A** Time sequence snapshots of microtubules growing from brightly-labeled seeds (magenta, Atto647) in the presence of 20 nM untagged KIF2A. Circles track the depolymerizing minus ends of selected microtubules that undergo treadmilling. A solid circle indicates the microtubule is still attached to the brightly labeled GMPCPP seed, whereas a dashed circle indicates the seed has completely depolymerized and the microtubule (including the minus end) is composed of only GTP/GDP tubulin. For better distinction and identification of the selected microtubules, both solid and dashed circles are color-coded. Arrows indicate the direction of “travel” across the glass surface. The asterisk indicates the microtubule was lost from view after this time-point following a plus-end catastrophe. **B** Representative kymographs of microtubules (magenta) that are released from the stabilized GMPCPP seed after depolymerization by mScarlet-KIF2A (cyan), indicated by the black arrowhead. Minus-end depolymerization was heterogeneous: asterisks indicate instances of increased depolymerization speed. Insets: enlarged regions of kymographs showing mScarlet-KIF2A localization at minus ends undergoing periods of “slow” (10 nM KIF2A inset) and “fast” (20 nM KIF2A inset) depolymerization. Inset scale bars: horizontal, 1 µm; vertical, 30 sec. **C** Average depolymerization rates of minus ends of treadmilling microtubules, measured from microtubules that are no longer attached to seeds, in the presence of various concentrations of mScarlet-KIF2A. Each measurement is plotted as a semi-transparent dot (with random x-jitter to improve visualization) and the mean is overlaid as a black bar. Inset: the same data with a 0 nM KIF2A timepoint included (where each measurement is made from each instance of depolymerization after catastrophe) and an extended y axis. Number of depolymerization speeds measured per condition: mScarlet-KIF2A: 0 nM, *n* = 264; 10 nM, *n* = 74; 20 nM, *n* = 76; 40 nM, *n* = 62; 60 nM, *n* = 68.

### γTuRC protects microtubule minus ends from KIF2A depolymerizing activity

Next, we wondered whether KIF2A can depolymerize the minus ends of microtubules that are nucleated by γTuRC, which typically nucleates microtubules in cells. We purified and immobilized human biotinylated and mBFP-tagged γTuRC on a functionalized glass surface via biotin-NeutrAvidin links, and observed by TIRF microscopy γTuRC-mediated microtubule nucleation in the presence of tubulin, as described previously (Consolati et al., 2020) (Fig. 4A). In the absence of mScarlet-KIF2A, microtubules nucleated with only the plus end growing and the minus end being tethered to an immobilized γTuRC (Fig. 4B, C – 0 nM KIF2A; Movie 4). Notably, after nucleation microtubules remained tethered to γTuRC throughout the entire duration of the experiment (Suppl. Fig. 3A), indicating that γTuRC stably caps minus ends after microtubule nucleation, in agreement with previous reports (Berman et al., 2023; Consolati et al., 2020; Rai et al., 2022; Thawani et al., 2020; Wieczorek et al., 2021).

**Figure 4.**
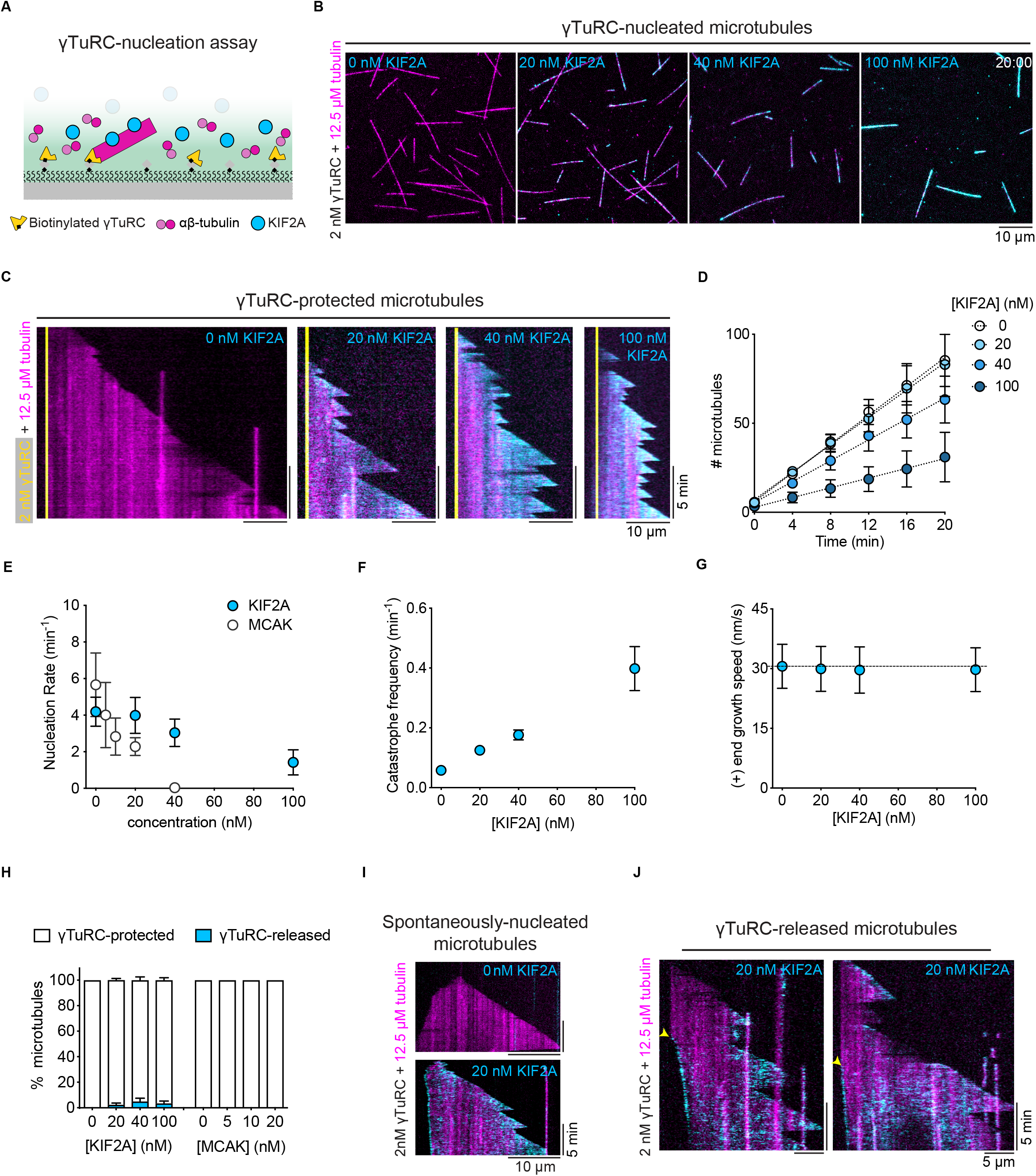
γTuRC-nucleated microtubules are protected from KIF2A depolymerizing activity. (**A**) Schematic of the TIRF microscopy-based γTuRC-nucleation assay. Biotinylated and mBFP-tagged γTuRC is immobilized on a biotin-PEG-functionalized glass surface via NeutrAvidin followed by the addition of tubulin and KIF2A. (**B)** TIRF microscopy images of γTuRC-nucleated microtubules in the absence (left) and presence of mScarlet-KIF2A (cyan) at the indicated concentrations and 12.5 μM of tubulin (AlexaFluor647, 5,4%; magenta), 20 minutes after the start of imaging. Surface-immobilized γTuRC is not shown. **(C)** Representative kymographs of microtubules nucleating from γTuRC (yellow) for the conditions shown in B. **(D)** Numbers of microtubules nucleated over time, **(E)** microtubule nucleation rates for mScarlet-KIF2A and mCherry-MCAK, **(F)** catastrophe frequencies and **(G)** microtubule plus-end growth speeds for the conditions in B. Circles represent mean values and error bars are SEM (D, E, F) and SD (G). For symbols without visible error bars, error bars are smaller than the symbol size. The lines in (D) represent linear regressions. The dashed line in (G) represents the mean microtubule plus-end grow speeds in the absence of mScarlet-KIF2A. Nucleation rates (E) were calculated from the slope of the linear regression (mScarlet-KIF2A: D; mCherry-MCAK: not shown). Number of microtubules analyzed for the catastrophe frequency (F) per mScarlet-KIF2A concentration: 0 nM, n = 191; 20 nM, n = 186; 40 nM, n = 122; 100 nM, n = 65; Number of microtubule growth speeds (G) measured per mScarlet-KIF2A concentration: 0 nM, n = 187; 20 nM, n = 169; 40 nM, n = 79; 100 nM, n = 66. Data for plots was pooled from three independent experiments. **(H)** Percentage of microtubules that either remain protected by γTuRC or are released after nucleation, for the indicated concentrations of mScarlet-KIF2A and mCherry-MCAK. Number of microtubules analyzed per condition: mScarlet-KIF2A – 0 nM, n = 256; 20 nM, n = 249; 40 nM, n = 190; 100 nM, n = 91; mCherry-MCAK – 0 nM, n = 332; 5 nM, n = 281; 10 nM, n = 182; 20 nM, n = 156. Data was pooled from three independent experiments. Error bars shown for mScarlet-KIF2A concentrations are SEM. **(I)** Representative kymographs of occasionally observed spontaneously nucleated microtubules, recognized by growth at both ends after nucleation, in a γTuRC nucleation assay, in the absence (top image) or presence (bottom image) of 20 nM of mScarlet-KIF2A (cyan). **(J)** Representative kymographs of γTuRC-nucleated microtubules that are released after nucleation in the presence of 20 nM of mScarlet-KIF2A (cyan). Yellow arrowheads indicate the moment of γTuRC-uncapping and the beginning of accumulation of mScarlet-KIF2A at the released and slowly depolymerizing microtubule minus end. 2 nM γTuRC was used for initial surface immobilization in all experiments.

Next, we added mScarlet-KIF2A at different concentrations to the γTuRC nucleation assay. Microtubules still nucleated from γTuRC, but nucleation became less efficient at higher KIF2A concentrations (Fig. 4B; Movie 4). Quantifying the number of nucleated microtubules over time revealed that the nucleation rate (slopes in Fig. 4D) decreased with increasing concentrations of mScarlet-KIF2A, but to a considerably lesser extent than with mCherry-MCAK (Fig. 4E), which was reported previously to inhibit γTuRC-mediated microtubule nucleation (Thawani et al., 2020). As observed for microtubules grown from GMPCPP-seeds (Fig. 2C), KIF2A at higher concentrations increased the catastrophe frequency of the plus ends of γTuRC-nucleated microtubules (Fig. 4C and F) without affecting their growth speed (Fig. 4G). These results indicate that plus end destabilization of the nascent microtubule nucleus forming on γTuRC negatively influences microtubule nucleation. Nevertheless, γTuRC-mediated nucleation tolerates much higher concentrations of KIF2A than MCAK, likely due to KIF2A’s end selectivity.

Remarkably, the large majority of the inherently more KIF2A-sensitive microtubule minus ends were now completely protected from the depolymerizing activity of KIF2A by the γTuRC cap (Fig. 4C). Despite the presence of KIF2A, almost all microtubule minus ends remained protected by γTuRC for the entire duration of the experiment, as demonstrated by their minus ends remaining tethered to the functionalized glass surface (Fig. 4C and H). This is in contrast to uncapped minus ends of dynamic microtubules (Fig. 2 and 3) or of occasionally spontaneously nucleated microtubules in the γTuRC-nucleation assay (Suppl. Fig. 3B), where the minus ends are depolymerized by KIF2A (Fig. 4I). Nevertheless, we still sporadically observed microtubules detaching from γTuRC in the presence of KIF2A (mScarlet-KIF2A: Fig. 4H; untagged KIF2A: Suppl. Fig. 3C). These microtubules first nucleated from γTuRC and remained minus end capped for a period of time until KIF2A apparently destabilized the connection with the γTuRC and drove slow minus end depolymerization (Fig. 4J; Movie 5). This was accompanied by minus end accumulation of KIF2A and resulted in treadmilling of the majority of γTuRC-released microtubules (Fig. 4J; Movie 5), as observed for seed-grown microtubules (Fig. 3B). In the absence of KIF2A or in the presence of mCherry-MCAK we did not observe a single released microtubule (Fig. 4H; Suppl. Fig. 3C), demonstrating that although microtubule release from γTuRC is a rare event, it can be selectively triggered by KIF2A.

### Together KIF2A and spastin uncap γTuRC-nucleated microtubules allowing microtubule treadmilling

This raises the question whether microtubule severases that have been reported to promote poleward microtubule flux in spindles during mitosis (Guerreiro et al., 2021; Huang et al., 2021; Zhang et al., 2007) can promote the generation of uncapped microtubules and therefore KIF2A-accessible minus ends of γTuRC-nucleated microtubules. To be able to test this, we purified human spastin and fluorescently labeled mGFP-spastin (Suppl. Fig. 1F, G) and verified their microtubule severing activity by adding them to glass-surface immobilized GMPCPP-stabilized microtubules. Although mGFP-spastin was a little less efficient than the unlabeled severase, both caused microtubule disintegration within minutes (Suppl. Fig. 4A, B – top panel), as it was observed for related severases (McNally et al., 1996; Roll-Mecak and Vale, 2005; Vemu et al., 2018). However, the vast majority of γTuRC-nucleated microtubules did not show any severing events in the presence of spastin (∼ 99%, Fig. 5A). This agrees with previous reports of microtubule severing being inefficient in the presence of free tubulin (Bailey et al., 2015; Jiang et al., 2017; Kuo et al., 2019; Vemu et al., 2018).

**Figure 5.**
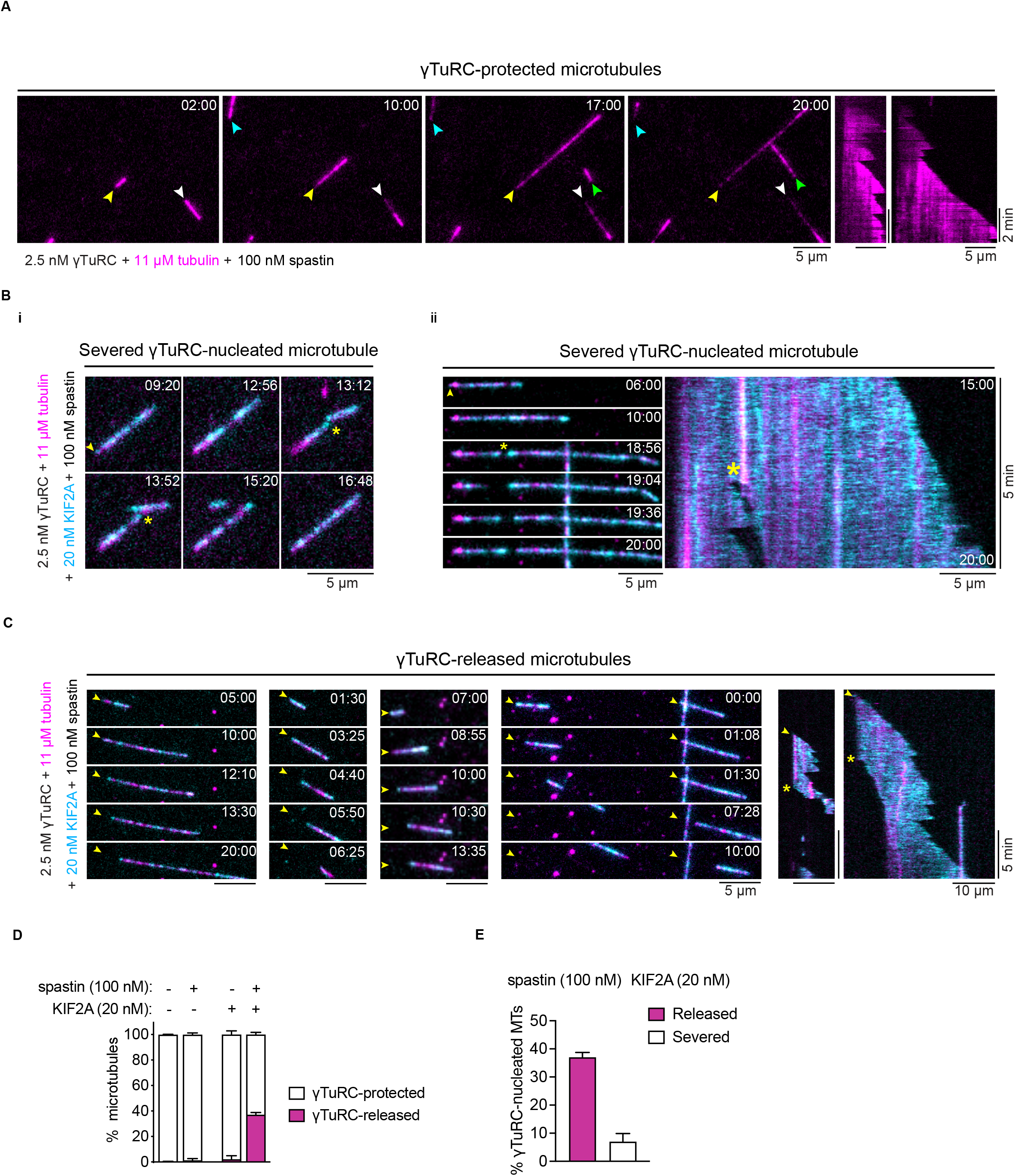
γTuRC-nucleated microtubules are uncapped by KIF2A and spastin allowing microtubule treadmilling. (**A**) Time sequence of TIRF microscopy images and kymographs of microtubules (magenta) nucleated by surface-immobilized γTuRC in the presence of 11 µM Cy5-tubulin (5.4% labelling percentage), which remain unaffected by 100 nM of spastin. Arrowheads point to γTuRC-capped microtubule minus ends. **(B)** Time sequence of TIRF microscopy images and kymograph of γTuRC-nucleated microtubules (magenta) that were severed in the presence of 11 µM Cy5-tubulin, 20 nM mScarlet-KIF2A (cyan) and 100 nM of spastin. Yellow arrowheads and asteriks indicate γTuRC-capped microtubule minus-ends and severing events, respectively. **(C)** Time sequence of TIRF microscopy images and kymographs of γTuRC-nucleated microtubules (magenta) that were released from γTuRC by the combined action of mScarlet-KIF2A (cyan) and spastin. Conditions as in (B). Yellow arrowheads point to the γTuRC-nucleation sites of microtubules and the yellow asterisks indicate the release of the microtubules from γTuRC, leaving the minus end uncapped for depolymerization by mScarlet-KIF2A (cyan). **(D)** Percentage of microtubules that either remain protected by γTuRC or are released after nucleation, for the indicated concentrations of mScarlet-KIF2A and spastin. Number of microtubules analyzed per condition: mScarlet-KIF2A and spastin – 0 nM, n = 228; mScarlet-KIF2A 0 nM and Spastin 100 nM, n = 152; mScarlet-KIF2A 20 nM and spastin 0 nM, n = 160; mScarlet-KIF2A 20 nM and spastin 100 nM, n = 222. Data for plots was pooled from at least two independent experiments. Error bars are SEM. **(E)** Percentage of released and severed γTuRC-nucleated microtubules for the indicated concentration of spastin and mScarlet-KIF2A. Number of microtubules analyzed per condition: mScarlet-KIF2A 20 nM and spastin 100 nM, n = 222. Error bars are SEM. In all experiments, 2.5 nM γTuRC was used for surface-immobilization. The time stamps indicate min:s.

When we added mGFP-spastin and mScarlet-KIF2A in combination to stabilized microtubules, they were now severed considerably faster than in the presence of spastin alone (Suppl. Fig. 4B – bottom panel). Measuring the fluorescence intensity along the initial microtubule contour over time revealed that when only GFP-spastin was present, its binding initially increased as the microtubule started to become severed and then decreased as the microtubule disappeared (Suppl. Fig. 4C). In the additional presence of KIF2A, this process exhibited a similar pattern but at an accelerated rate (Suppl. Fig. 4C), suggesting that KIF2A supports severing by spastin and that both act in synergy. Indeed, KIF2A’s fluorescence intensity peaked right after the peak of the mGFP-spastin intensity (Suppl. Fig. 4C).

When we added both spastin and KIF2A to the γTuRC nucleation assay, we also observed an increased number of severing events along the lattice of γTuRC-nucleated microtubules compared to spastin being present alone (yellow asterisks, Fig. 5B i and ii; severing observed in ∼7 % of microtubules). After severing, the newly formed plus ends displayed dynamic instability behavior switching between growth and fast shrinkage episodes (Fig. 5B; Movie 6), whereas the newly formed minus ends were slowly depolymerized by KIF2A (Fig. 5B), resulting in treadmilling of the newly generated free microtubule (Fig. 5B ii). Remarkably, in the combined presence of mScarlet-KIF2A and spastin we observed also substantially more frequent detachments of microtubules from surface-tethered γTuRC, also causing them to treadmill (Fig. 5C; Movie 7). Microtubule release from immoblized γTuRC increased from ∼1 % and ∼2 % in the presence of spastin or KIF2A alone, respectively, to almost 40% when spastin and KIF2A were both present (Fig. 5D). This release, induced by severing at or near the γTuRC/microtubule interface, is ∼5 times higher than the percentage of microtubules severed at any other point along the lattice (Fig. 5E), despite the much larger number of potential severing sites along the entire microtubule length compared to the single γTuRC cap at the minus end. Because we did not observe mGFP-spastin directly binding to surface-immobilized γTuRC (Suppl. Fig. 4D), we conclude that the γTuRC/microtubule interface is orders of magnitudes more sensitive to the combined action of KIF2A and spastin than an average microtubule lattice site, favoring γTuRC uncapping over lattice severing. Taken together, our results demonstrate that spastin and KIF2A act in synergy to trigger microtubule minus-end release from γTuRC and subsequent treadmilling, consistent with the proposed function of KIF2A and severases at spindle poles.

## DISCUSSION

We discovered here that the kinesin-13 KIF2A is a microtubule minus-end depolymerase that can induce flux-like microtubule treadmilling. Although KIF2A acts mildly on plus ends, it has a strong preference for minus end destabilization, establishing it as an asymmetric microtubule depolymerase. This property distinguishes KIF2A from its paralog MCAK that destabilizes both microtubule ends similarly. Moreover, we found that the presence of a γTuRC cap at minus ends inhibits minus-end depolymerization by both depolymerases. To induce treadmilling of γTuRC-nucleated microtubules, active uncapping of minus ends is required, which can be achieved by the combined activities of KIF2A and the severase spastin. Taken together, we show here that a γTuRC/KIF2A/severase module can provide the activities required for both microtubule nucleation and controlled microtubule minus-end depolymerization at spindle poles, in agreement with the requirements for the generation of microtubule flux in mitotic and meiotic spindles.

Minus end recognition by KIF2A appears to be different from the generic microtubule end recognition mechanism of kinesin-13s, in which the motor domain in its ATP state binds to and stabilizes a curved microtubule protofilament (Asenjo et al., 2013; Benoit et al., 2018; Trofimova et al., 2018; Wang et al., 2017). We found that KIF2A binds preferentially to minus ends also in the presence of ADP, showing that enzymatic activity is not required for minus end recognition. Because ADP-KIF2A also slows down microtubule growth and because minus-end capping by iE5 and γTuRC prevents KIF2A minus-end accumulation, it is tempting to speculate that KIF2A’s minus end recognition may involve binding to at least part of the exposed α-tubulin at microtubule minus ends. In the future, biochemical and structural studies will be required to identify the mechanism underlying KIF2A’s remarkable minus end binding preference.

In interphase cells, individual microtubules have been observed to adopt a treadmilling state in which net plus end growth is balanced by net minus end depolymerization, which has been associated with loss of minus-end stabilizers (Goodwin and Vale, 2010; Rodionov and Borisy, 1997; Shaw et al., 2003; Waterman-Storer and Salmon, 1997). Remarkably, the kinesin-13 KLP10A, which in mitosis seems to share some functional homology with KIF2A (Laycock et al., 2006; Rath et al., 2009; Rogers et al., 2004), was observed to localize to shrinking minus ends and to induce treadmilling of microtubules in *Drosophila* cells when the minus-end stabilizing protein patronin was knocked down (Goodwin and Vale, 2010).

Early *in vitro* work with purified proteins investigated the possibility of treadmilling being an intrinsic property of microtubules, but later it became clear that the intrinsic dynamic behavior of microtubules is dominated by dynamic instability at both ends (Grego et al., 2001; Margolis and Wilson, 1978; Mitchison and Kirschner, 1984). We find here that purified KIF2A can trigger microtubule treadmilling *in vitro*, resulting as the consequence of the combination of two of its properties. First, it destabilizes minus ends more strongly than plus ends, so that in a certain concentration regime plus ends grow dynamically whereas minus end shrink; and second, in contrast to the intrinsically fast depolymerization after typical catastrophe events, KIF2A induces mainly slow minus end depolymerization, while remaining end-accumulated at the shrinking minus ends, which can then be balanced by (net) plus end growth. The mechanism of this slow-down of minus end depolymerization is unknown, but might be related to the formation of higher order structures formed by curved, depolymerizing protofilaments and KIF2A, as observed in electron microscopy with different members of the kinesin-13 family (Moores et al., 2006; Tan et al., 2006; Tan et al., 2008; Zhang et al., 2013).

A recent study reported the *in vitro* reconstitution of microtubule treadmilling with a combination of several vertebrate microtubule binding proteins (Arpağ et al., 2020), evidence of which can also be seen in earlier *in vitro* work with the corresponding *Drosophila* orthologs (Moriwaki and Goshima, 2016). Because the symmetric depolymerase MCAK was used in this recent work, additional activities were required to generate an asymmetry in the control of microtubule dynamics. This was achieved by including the plus-end specific microtubule polymerase XMAP215, and the anti-catastrophe factor CLASP2, resulting in microtubule treadmilling with growing plus ends and shrinking minus ends. These additional factors were not required in our reconstitution, due to the inherently asymmetric activity of KIF2A with respect to the two microtubule ends, simplifying the minimal requirements for microtubule treadmilling. In cells KIF2A’s intrinsic minus end preference can be enhanced by proteins that promote its minus end accumulation (Guan et al., 2023).

Our study revealed that γTuRC and kinesin-13s compete in two different ways. First, we showed that microtubule nucleation by γTuRC is inhibited in a dose-dependent manner by both KIF2A and MCAK, however to a much lesser extent by KIF2A. This suggests that kinesin-13 depolymerase activity destabilizes nascent microtubule plus ends beginning to grow on a γTuRC template, and that this destabilization is weaker for KIF2A due to its end selectivity. Second, once the microtubule has nucleated, γTuRC caps the minus end and now inhibits minus end depolymerization by kinesin-13s, an activity reminiscent of the minus-end binding protein patronin/CAMSAP that however binds minus ends very differently from γTuRC (Atherton et al., 2017; Goodwin and Vale, 2010).

Our finding that spastin and KIF2A in combination, but not alone, can efficiently induce microtubule severing in the presence of free tubulin leading then to KIF2A-mediated minus end depolymerization has several interesting implications. First, severase-induced lattice damage appears to provide an entry point for KIF2A to promote lattice destabilization, and/or interfere with lattice repair by soluble tubulin (Schaedel et al., 2015; Vemu et al., 2018), thereby promoting complete severing. KIF2A’s severase-supporting effect may explain why in cell extract severases appear much more active than purified severases in the presence of free tubulin *in vitro* (Vale, 1991; Vemu et al., 2018). α-tubulins likely become exposed when spastin removes tubulins from the lattice where KIF2A might be able to act and increase the size of the damage site. Our results also highlight that severase activity in the presence of free tubulin can be promoted by mechanisms beyond physical recruitment of severases to the microtubule, as shown for ASPM and katanin (Jiang et al., 2017).

Moreover, we observed that also γTuRC-uncapping was promoted by the combined action of KIF2A and spastin, and remarkably was more frequent than microtubule severing, although (i) γTuRC prevents microtubule minus end binding of KIF2A, (ii) γTuRC and spastin do not directly interact, and (iii) many more potential severing sites exist along the microtubule lattice than at the γTuRC/microtubule minus end interface. This implies that the γTuRC/minus end connection is much more sensitive to the combined action of spastin and KIF2A than the rest of the microtubule lattice. Although the γTuRC cap at minus ends is very stable in the absence of other proteins, it appears to become a structurally weak point in the presence of KIF2A and a severase, allowing preferential active minus-end uncapping, possibly not only to enable microtubule treadmilling, but also to recycle γTuRC for new rounds of microtubule nucleation. γTuRC-uncapping by KIF2A and spastin is functionally distinct from recently reported uncapping by CAMSAPs that remove γTuRC by inducing slow minus end polymerization, which is possibly more adapted to the function of microtubules in neurons (Rai et al., 2022).

In conclusion, we characterized *in vitro* a minimal protein module that can be found at the poles of mitotic and meiotic spindles during cell division and that can control apparently antagonistic activities at this location. On one hand, microtubules need to be nucleated which results in stable capping of their minus ends by γTuRC. On the other hand, minus ends also need to be uncapped and slowly shrink for microtubule flux in the spindle to occur. γTuRC, KIF2A and a severase in combination can provide this control of microtubule minus end dynamics as required at spindle poles. A major element of this control is the asymmetric action of the kinesin-13 KIF2A, adding to the list of other plus or minus end selective microtubule associated proteins that control the functionally important asymmetric dynamic properties of the two microtubule ends in cells.

## ACKNOWLEDGEMENTS

We thank Benoit Gigant for providing the αRep iE5 plasmid, Felix Ruhnow for microscopy and data analysis support, Maria Gili for insect cell expression of KIF2A constructs, Johanna Roostalu for the design of the MCAK and spastin expression constructs, Raquel Garcia-Castellanos for technical support, the Cell Services of the Francis Crick Institute for producing large HeLa cell cultures. This work was supported by the Spanish Ministry of Science and Innovation to the EMBL partnership, the Centro de Excelencia Severo Ochoa and the CERCA Programme of the Generalitat de Catalunya, and by the Francis Crick Institute, which receives its core funding from Cancer Research UK (FC001163), the UK Medical Research Council (FC001163), and the Wellcome Trust (FC001163). C.B. was supported by EMBO longterm fellowship ALTF-883-2020 and Marie Curie fellowship TuRCReg. T.S. acknowledges support from the European Research Council (ERC Synergy Grant, Project 951430) and from the Spanish Ministry of Science and Innovation (grant PID2019-108415GB-I00).

## AUTHOR CONTRIBUTIONS

G.H., C.B., and T.S. designed research; G.H., C.B. and C.T. generated reagents, G.H. and C.B. performed all other experiments and analyzed data; G.H., C.B., and T.S. wrote the paper.

## METHODS

### Protein constructs and cloning

Human KIF2A (NCBI reference sequence: NP_004511.2; codon-optimized for insect cell expression by GeneArt) and MCAK (NCBI reference sequence: NP_006836.2) constructs were cloned similarly to previous published constructs (Roostalu et al., 2018). Briefly, coding sequences were amplified by PCR and cloned into a pFastBac expression vector for expression in *Sf*9 or *Sf*21 cells (Thermo Fisher), including a StrepTagII-coding sequence at the N terminal of the construct, to generate the following constructs: StrepTagII-KIF2A, StrepTagII-mScarlet-G_6_S-KIF2A and StrepTagII-mCherry-G_5_A-MCAK. Constructs included a region coding for a TEV protease recognition site between the StrepTagII sequence and the desired construct for cleavage of the tag. The monomeric red fluorescent protein variants mScarlet (Bindels et al., 2017) and mCherry (Shaner et al., 2004) were separated from the N terminus of KIF2A and MCAK by flexible linker regions. The sequence coding for MCAK included an extra isoleucine residue after the initial methionine as a consequence of the cloning process.

The human spastin and mGFP-spastin constructs were cloned using the sequence corresponding to isoform 3 (UniProtKB reference sequence Q9UBP0-3; OriGene), a shortened variant derived from use of an alternative start residue (Met-87), as identified and used in previous work (Claudiani et al., 2005; Roll-Mecak and Vale, 2005). The spastin sequence was amplified by PCR and cloned into a pETMZ and a pETMZ-mGFP vector for expression in *E. coli*, generating the constructs His_6_-Ztag-TEV-spastin and His_6_-Ztag-TEV-mGFP-spastin. The sequence coding for spastin included an extra isoleucine residue after the initial methionine as a consequence of the cloning process.

The plasmids for bacterial expression of the αRep iE5 and the tandem repeat Design-Ankirin-Repeat-Protein (DARPin) – (D1)_2_ – are described elsewhere (Campanacci et al., 2019; Pecqueur et al., 2012).

### Protein expression and purification

Tubulin was prepared from pig brains as previously described (Castoldi and Popov, 2003). Tubulin was further purified by recycling and some tubulin was labeled with Alexa647-N-hydroxysuccinimide ester (NHS; Sigma-Aldrich), Atto647-NHS (Sigma-Aldrich), Cy5-NHS (Lumiprobe), or biotin-NHS (Thermo Scientific), as described (Consolati et al., 2022). Final concentrations and fluorescent labeling ratios were determined by UV absorption using NanoDrop™ One/OneC (Thermo Scientific) (extinction coefficient for tubulin, 115,000 M^−1^ cm^−1^; molecular weight, 110 kDa) after final ultracentrifugation, before aliquoting and snap-freezing. Aliquoted tubulin was then in liquid nitrogen.

KIF2A, mScarlet-KIF2A and mCherry-MCAK constructs were expressed using *Sf*21 cells (Thermo Fisher) according to the manufacturer’s protocols. Cell pellets from 500 mL of culture were snap-frozen and stored at –80°C. Thawed pellets of cells expressing KIF2A and mCherry-MCAK constructs were resuspended in ice-cold KIF2A lysis buffer (50 mM Na-phosphate, 300 mM KCl, 1 mM MgCl_2_, 1 mM ethylene glycol bis(2-aminoethyl ether)tetraacetic acid (EGTA), 5 mM β-mercaptoethanol (β-ME), and 0.5 mM adenosine triphosphate (ATP), pH 7.5), supplemented with DNaseI (Sigma-Aldrich) and protease inhibitors (cOmplete EDTA-free, Roche). The lysate was homogenized in a glass douncer on ice (40 strokes) and subsequently clarified by centrifugation at 50,000 rpm in a Ti70 rotor (Beckman Coulter) at 4°C for 45 min. The supernatant was then run over a 5 mL StrepTrap HP or XT column (Cytiva) at 0.5 mL min^-1^ using a peristaltic pump at 4°C. Bound protein was washed with 10 mL of KIF2A lysis buffer supplemented with ATP to a total of 5 mM at 0.5 mL min^-1^ and then eluted in 0.5 mL fractions using KIF2A lysis buffer supplemented with 2.5 mM d-desthiobiotin (for StrepTrap HP) or 50 mM biotin (for StrepTrap XT). Protein-containing fractions were collected, pooled, supplemented with protease inhibitors, and cleaved with TEV protease overnight on ice. The buffer was then exchanged to KIF2A gel filtration buffer (lysis buffer with 0.1 mM ATP instead of 0.5 mM) using HiPrep 26/10 or PD10 desalting columns (Cytiva) to remove the d-desthiobiotin/biotin, and the protein was run over the StrepTrap column again at 0.5 mL min^-1^ and 4°C to obtain only cleaved protein. The cleaved protein was then concentrated (Vivaspin 30,000 MWCO, Sartorius) and ultracentrifuged in a TLA-120.2 rotor (Beckman Coulter) at 80,000 rpm for 10 minutes at 4°C before further purification by gel filtration using a HiLoad 16/600 Superose 6 prep grade column (Cytiva), equilibrated in KIF2A gel filtration buffer. Protein eluted as a single peak which was collected and concentrated to 3-5 mg mL^-1^, ultracentrifuged in a TLA 120.2 rotor (Beckman Coulter) at 80,000 rpm for 15 minutes at 4°C, snap-frozen and stored in liquid nitrogen. mCherry-MCAK was supplemented with 10% sucrose (wt/vol) before the final ultracentrifugation.

mScarlet-KIF2A was purified similarly to KIF2A and mCherry-MCAK, with the following modifications: the lysate was homogenized using the EmulsiFlex-C5 (10,000 – 15,000 psi, Avestin). Affinity chromatography was performed using the Äkta Pure^TM^ System (Cytiva) instead of the peristaltic pump and gel filtration using a HiLoad 16/600 Superdex 200 pg column (Cytiva). mScarlet-KIF2A was also supplemented with 10% sucrose (wt/vol) before the final ultracentrifugation.

His_6_-Ztag-spastin was expressed in BL21 pRIL *E. coli* cultures at 18°C overnight, pellets of which were snap-frozen and stored at –80°C. For purification, pellets were resuspended in spastin lysis buffer (50 mM 4-(2-hydroxyethyl)-1-piperazineethanesulfonic acid (HEPES), 300 mM KCl, 5 mM MgCl_2_, 25 mM sucrose, 0.5 mM ATP, 1 mM β-ME, pH 8.0) supplemented with DNAse I and protease inhibitors. Cells were lysed using a microfluidizer and cleared by centrifugation. Cleared lysate was loaded onto a HiTrap column loaded with cobalt (Cytiva) and washed with spastin wash buffer (50 mM HEPES, 300 mM KCl, 2 mM MgCl_2_, 1 mM β-ME, 50 mM arginine, 50 mM glutamate, 25 mM sucrose, 0.1 mM ATP, 20 mM imidazole, pH 7.5) before eluting with spastin wash buffer supplemented with imidazole to a total of 400 mM. Eluted protein fractions were pooled and the buffer was exchanged with spastin wash buffer using PD-10 columns and cleaved with TEV protease overnight. Uncleaved protein and cleaved His_6_-Ztag domains were separated from cleaved protein by incubation with TALON resin (Clontech). The supernatant was buffer exchanged to MES A buffer (20 mM MES, 100 mM KCl, 2 mM MgCl_2_, 0.1 mM ATP, 5 mM β-ME, pH 6.0) using PD-10 columns. The buffer-exchanged protein was ultracentrifuged and bound to a Mono S 5/50 GL ion exchange column equilibrated in MES A. Protein was eluted from the column with a gradient of MES B buffer (20 mM MES, 1000 mM KCl, 2 mM MgC_2_, 0.1 mM ATP, 5 mM β-ME, pH 6.0; 3% mL^-1^ gradient). Eluted protein was then gel-filtered using a HiLoad 16/600 Superdex 200 pg column (Cytiva) equilibrated with spastin gel-filtration buffer (50 mM HEPES, 300 mM KCl, 0.1 mM ATP, 2 mM MgCl_2_, 5 mM β-ME, 50 mM arginine, 50 mM glutamate, 250 mM sucrose, pH 7.5). Pooled fractions were then concentrated, ultracentrifuged, aliquoted and snap-frozen, and stored in liquid nitrogen. Before use in experiments, frozen aliquots were thawed and dialyzed against KIF2A gel-filtration buffer (Slide-A-Lyzer MINI, Thermo Fisher). Dialyzed protein was then concentrated, ultra-centrifuged in a TLA 120.2 rotor at 80,000 rpm for 10 minutes at 4°C, snap-frozen, and stored in liquid nitrogen.

His_6_-Ztag-mGFP-spastin was expressed and purified using the same procedure as for human spastin.

KIF2A, mScarlet-KIF2A, mCherry-MCAK and spastin concentrations were all measured by Bradford assay (Bio-Rad) against a bovine serum albumin standard (Pierce/Thermo Fisher) using snap-frozen and thawed protein aliquots. Reported concentrations refer to monomers based on their predicted molecular weights.

Biotinylated γTuRC-mBFP-AviTag was purified from HeLa-Kyoto cells stably expressing GCP2-mBFP-AviTag and the biotin ligase HA-BirA, essentially as previously described (Consolati et al., 2020). γTuRC-mBFP-AviTag concentration was estimated by UV absorption using NanoDrop™ One/OneC (Thermo Scientific) (MW: 2.2 MDa), yielding 0.4-0.8 mg from 60 g of HeLa-Kyoto cells.

The αRep iE5 was expressed and purified as described elsewhere (Campanacci et al., 2019).

The DARPin (D1)_2_ (Pecqueur et al., 2012) was expressed in *E. coli* BL21-CodonPlus (DE3)-RIL cells grown in LB medium. When reaching an OD_600_ of 0.6 at 37°C, expression of (D1)_2_ was induced with 0.5 mM isopropyl β-D-1-thiogalactopyranoside (IPTG) for 20 hours at 18°C. Cells from 500 mL expressing cultures were harvested by centrifugation (9,000 x g for 10 minutes, at 4°C), and cell pellets were stored at –80°C. (D1)_2_ was purified as previously described (Pecqueur et al., 2012), with some modifications. Briefly, cells were resuspended in lysis buffer (50 mM Tris, pH 8, 1 mM MgCl_2_, 10 mM imidazole; 10 mL per gram of pellet) and supplemented with 0.3 mg mL^-1^ lysozyme, 0.25 units mL^-1^ of benzonase and protease inhibitor mix (cOmplete EDTA free, Roche). Cells were lysed using the EmulsiFlex-C5 (15,000 psi, Avestin) and lysate was clarified by centrifugation at 20,000 × g for 40 minutes at 4°C. The supernatant was loaded using a peristaltic pump in 1 mL (bed volume) of Ni-NTA agarose (QIAGEN) packed in a Poly-Prep® Chromatography Column (Cat #731-1550, BioRad). Bound protein was washed with 30 mL of wash buffer (20 mM K-phosphate, pH 7.2, 1 mM MgCl_2_, 0.5 mM EGTA, 100 mM KCl, 1 mM ATP, 10 mM imidazole and 5 mM DTT), followed by elution in wash buffer without ATP and 100 mM imidazole instead of 10 mM. Eluted protein was subjected to size exclusion chromatography on a HiLoad 16/600 Superdex 200 pg column (Cytiva) equilibrated with wash buffer without ATP and imidazole. Protein-containing fractions were pooled, concentrated using an Amicon centrifugal unit (MWCO 10,000, Millipore) and ultracentrifuged in a TLA110 rotor (Beckman Coulter) at 80,000 rpm for 10 minutes at 4°C.

(D1)_2_ and iE5 concentrations were estimated by UV absorption using NanoDrop™ One/OneC (Thermo Scientific) (extinction coefficients: 13,980 M^-1^ cm^-1^ and 5,960 M^-1^ cm^-1^; molecular weights: 35.2 kDa and 25.1 kDa, respectively).

### Glass

Polyethylene glycol (PEG)-passivated glass, functionalized with biotin, was prepared as described previously (Consolati et al., 2022). Glass for γTuRC nucleation assays was generally used within two weeks; otherwise, glass was generally used within eight weeks.

### Stabilized GMPCPP seeds

Stabilized microtubules (seeds) were polymerized using the slowly-hydrolyzing GTP analog, GMPCPP (Jena Bioscience). Seeds were polymerized from 3 µM of tubulin in BRB80 (80 mM K-PIPES, 1 mM MgCl_2_, 1 mM EGTA, pH 6.8; prepared and stored as a 5× stock for up to 2 weeks), including 18 % biotin-labeled tubulin, as well as fluorescent tubulin (AlexaFluor674-NHS or Atto647-NHS) for a final fluorescent labeling ratio of 3-5% for “dim” seeds (used for measurements of seed depolymerization in Figure 1) or 10-15% for “bright” seeds (used for dynamic assays in Figures 2 and 3). 200 µL of the tubulin mixture, including 0.5 mM GMPCPP, was incubated on ice for 5 minutes before transferring to a heat block at 37°C for one hour to initiate nucleation and elongation. The microtubules were then diluted with 170 µL warm BRB80 and centrifuged at 17,000 x g for 10 minutes in a tabletop centrifuge. The microtubule pellet was washed by the addition of 400 µL warm BRB80 and subsequent centrifugation for another 10 minutes at 17,000 x g in the tabletop centrifuge. The washed microtubule pellet was finally resuspended in 50 µL warm BRB80. The microtubules were further diluted 100 – 300 fold in BRB80 at room temperature before microscopy experiments. The microtubules were generally used on the same day. 3 µM of total tubulin resulted in relatively long microtubules, allowing for longer observation times of depolymerization activities, but could be sheared by 10 passes through a 27-gauge needle to generate shorter seeds, for shorter seed depolymerization times and therefore more treadmilling events within the experimental timeframe (Fig. 3).

### Polarity-marked seeds

To grow microtubules with polarity-marked segments with both segments being stabilized by GMPCPP, we developed a new protocol using the microtubule plus end capping DARPin (D1)_2_ (Pecqueur et al., 2012). Firstly, dim seeds (3-5% label ratio, 18% biotin tubulin) were first prepared as above. Subsequently, a bright GMPCPP mix (10-15% label ratio, 18% biotin tubulin) was prepared on ice, as above. While incubating the mix on ice, 2 µL of 2.5-5 µM DARPin (D1)_2_ (diluted in BRB80) was added to 8 µL of the dim seeds. The bright mix was then diluted 1:1 (to 1.5 µM tubulin, 0.25 mM GMPCPP) and 90 µL of the diluted bright mix was allowed to warm for 60 seconds at 37°C before adding the 10 µL DARPin (D1)_2_ /dim seed mixture, resulting in a final DARPin (D1)_2_ concentration of 50-100 nM. Bright extensions were allowed to polymerize by incubation for one hour at 37°C before diluting, spinning, washing, spinning, and resuspending as above, except seeds were diluted 1 in 10 or 1 in 6 from the 50 µL resuspension before experiments.

Correct polarity-marking of the marked microtubules was confirmed using a gliding assay: flow chambers were assembled from a poly-(L-lysine)-PEG (SuSoS)-passivated counter glass and an untreated cover glass separated by double sticky tape. The chamber was washed with 50 µL gliding buffer (BRB80 with 5 mM β-ME, 1 mM ATP, 1 mg mL^-1^ β-casein), using filter paper to pull solution through the chamber, and incubated for 3 minutes. Then the chamber was washed with 50 µL of gliding buffer supplemented with 100 nM of the minus-end-directed motor mCherry-HSET (purified as described (Roostalu et al., 2018)), and incubated further 3 minutes. Lastly, 50 µL of room-temperature gliding buffer supplemented with polarity marked seeds was flowed into the chamber. Subsequently the chamber was sealed with silicone vacuum grease and imaged on a confocal microscope. 100% of motile microtubules with one bright and one dim segment moved in the direction of their dim end (*n* = 37 microtubules across 2 experiments), indicating that the bright end is the minus end and that our method of marking the ends is effective.

### TIRF microscopy experiments

KIF2A gel filtration buffer (without ATP or β-ME) and 5× BRB80 were prepared and stored at 4°C for a maximum of 2 weeks, and supplemented with ATP and β-ME, or diluted appropriately in MilliQ water, on each day of experiments. Buffers were filtered (0.22 µm pore size filters) or ultracentrifuged (80,000 rpm, 10 minutes, 4°C) and stored on ice for the day (Consolati et al., 2020). Flow chambers were assembled from poly-(L-lysine)-PEG-passivated counter glass, biotin-PEG functionalized cover glass, and double-stick tape, as described elsewhere (Consolati et al., 2022).

The assembled flow chamber was incubated for 10 minutes with 50 µL 5% Pluronic F-127 (Sigma Aldrich) in MilliQ water followed by two washes with 50 µL of casein buffer (0.2 mg mL^-1^ κ-casein (Sigma Aldrich), diluted in BRB80 or the assay buffer). The chamber was subsequently incubated for 5 minutes with 50 µL of 50 µg mL^-1^ of NeutrAvidin (Life Technologies) in casein buffer, followed by 2 washes with 50 µL of room-temperature BRB80 (seed assays; Figures 1-3) or with 50 µL of cold γTuRC buffer (γTuRC nucleation assays; Figures 4 and 5). The chamber was then incubated with 50 µL of biotinylated GMPCPP seeds or 50 µL of γTuRC pre-diluted in γTuRC buffer to the concentration indicated for each experiment. GMPCPP seeds or γTuRC were allowed to bind to the NeutrAvidin surface for 3-5 minutes followed by 2 washes with 50 µL of BRB80-based assay buffer (80 mM PIPES, 60 mM KCl, 5 mM Na-phosphate, 1.1 mM EGTA, 2 mM MgCl_2_, 1 mM GTP, 1 mM ATP (pH 7; alternatively, ADP or AMPPNP, pH 7), 5 mM β-ME, 0.15% (w/vol) methylcellulose (4,000 cP, Sigma-Aldrich) 1% (w/vol) glucose, 0.02% (vol/vol) Brij-35). The final experimental mix was then flowed in two steps of 25 µL, waiting 30 – 60 seconds between flows, to ensure equilibration of the chamber, and finally sealed with silicone vacuum grease (Obermeier). For the treadmilling experiments, the final assay mix was flowed in on a 33°C heat block to promote growth from small seed fragments in the presence of depolymerase activity. The final experimental mix was prepared on ice: assay buffer supplemented with oxygen scavengers (0.1 mg mL^-1^ catalase (Sigma-Aldrich), 1 mg mL^-1^ glucose oxidase (Serva)), either no tubulin (for ‘seed only’ and ‘γTuRC – mGFP-spastin co-localization’ experiments) or 11-22.5 µM of tubulin (containing 3-5% AlexaFluor647, Atto647, or Cy5-labelled tubulin, as indicated, for experiments with dynamically growing microtubules), the different enzymes (KIF2A and mScarlet-KIF2A, 0.5-250 nM, mCherry-MCAK, 5-100 nM, spastin 50 and 100 nM, and/or mGFP-spastin 50 and 100 nM as indicated) and the microtubule end capping proteins (iE5, 2 and 10 µM, and/or (D1)_2_, 3 and 15 µM). Their concentrations were altered by predilution in KIF2A gel filtration buffer (for experiments using 1 mM ADP or AMPPNP, KIF2A gel-filtration buffer was prepared with the corresponding nucleotides for diluting the proteins such that residual ATP from the storage buffer amounted to less than 0.0008 mM). KIF2A, mScarlet-KIF2A, mCherry-MCAK, iE5, (D1)_2_, spastin and/or mGFP-spastin dilutions in KIF2A gel-filtration buffer accounted for 10% of the final mix volume; otherwise, the equivalent volume of KIF2A gel-filtration buffer was used instead. The experimental mix was finally centrifuged in a 4°C tabletop centrifuge at 17,000 x g for 5 minutes, and the supernatant was recovered and returned to a tube on ice.

Polymerization from the seeds or nucleation from γTuRC was initiated with a temperature shift to 33°C in the TIRF microscope incubator (Okolab) and imaging was started within 2 minutes from the initial temperature shift. For ‘γTuRC – mGFP-spastin co-localization’ experiments, images were acquired 5 minutes after shifting the temperature to 33°C on the TIRF microscope.

### TIRF microscopy

Experiments were performed on a TIRF microscope (Cairn Research, Faversham, UK) (Hannabuss et al., 2019) using a 100× 1.49 NA Nikon Objective lens. Time-lapse imaging in multiple colors was performed between 1 frame/ 5 s and 1 frame / 2 s. γTuRC-mBFP-AviTag was imaged every frame (Fig. 4B, C, I and J; Fig. 5A; Movies 4 and 5) or once every 30 frames (Fig. 5B and C; Movies 6 and 7). For the ‘γTuRC – mGFP-spastin co-localization’ experiment (Suppl. Fig. 4D) stream acquisition of 100 frames in 10 seconds was performed sequentially for each fluorescence channel. Images were acquired with 300 ms exposure times for all channels (γTuRC-mBFP-AviTag; 405 nm excitation; mGFP-spastin; 488 nm excitation; mScarlet-KIF2A/mCherry-MCAK, 561 nm; Atto647/AlexaFluor647/Cy5-tubulin, 638 nm). Experiments were observed for 20-30 minutes. 2 or 3 independent experiments were performed per condition.

### TIRF microscopy image processing

Unless otherwise stated, all image processing and analysis was performed using Fiji (Schindelin et al., 2012). Recorded movies were first drift-corrected using the Descriptor-based stack registration plug-in, developed by Stephan Preibisch and included with Fiji, using the tubulin channel as the registration channel, and further background-subtracted (rolling ball; 50 pixels) prior to further analysis. To measure speeds and intensities, kymographs (space-time plots) were generated from drift-corrected videos using Fiji. For the γTuRC single-molecule visualization shown in Figure 4C, Supplementary Figure 3A, and Movie 5, an average γTuRC image was generated using the “Z project” function in Fiji. The average projection was then used as a static background merged with the other channels. For the ‘γTuRC – mGFP-spastin co-localization’ experiment (Suppl. Fig. 4D), γTuRC and mGFP-spastin images were averaged using the “Z project” function in Fiji. Tracking arrowheads in Movies 3, 5 and 7 were generated using a published plug-in (Daetwyler et al., 2020).

### Seed depolymerization speed

From kymographs of each microtubule, depolymerization speeds were measured manually by drawing a line along the shrinking ends, beginning with the first frame and ending with the last frame before the microtubule detaches or disappears, or the last frame of the video. The speed was calculated from the slope of the line. The shrinking could be directly attributed to minus or plus ends in the case of polarity-marked microtubules, which had brighter minus ends. For the KIF2A concentration series data (Fig. 1E) the polarity of the microtubule was inferred by the relative speed of the shrinking ends.

### End intensity measurement

Kymographs for end intensity measurement were cropped to 50 pixels in height (corresponding to 50 frames over 245 seconds). A segmented line tracing each end of polarity-marked microtubules (visualized in the tubulin channel) was drawn on the kymograph and used to measure the average intensity of mScarlet-KIF2A or mCherry-MCAK localizing to the ends (measured in the mScarlet/mCherry channel). The same measurement was made 6 pixels outside of the microtubule area, to account for local background, and 6 pixels inside the microtubule area of the kymograph, to measure the intensity on the lattice. Microtubules to which a large aggregate (>5x the average end intensity) of mScarlet-KIF2A or mCherry-MCAK bound within measured pixels were excluded from the analysis, and represented a minority of the cases. Intensity measurements for each end and the corresponding lattice measurements were corrected by subtracting the background measured from that end. The “per end” lattice intensities were then averaged to yield an average lattice intensity representing the whole microtubule, and then used to calculate the end intensity ratios. The “raw” intensities (Fig. 1H) were pooled from experiments done on different days with different laser settings and the average lattice intensity of mScarlet-KIF2A in AMPPNP, also measured on each day, was used to normalize the values.

### Microtubule dynamics

Kymographs of dynamic microtubules were used to determine the speeds of growing and shrinking phases, measured and calculated similarly to the depolymerization rates of seeds described above. Only measurements with a non-zero slope (and therefore measurable speed) were included for analysis. Microtubule polarity of dynamic microtubules grown from stabilized seeds was determined by comparison of the relative speeds of growth of the ends and comparison to the growth speeds of control microtubules (growing in the absence of added KIF2A, spastin or end cappers). For microtubules grown from seeds, the catastrophe frequencies of individual microtubules were calculated as the number of catastrophe events (switching from growth to shrinkage) detected in a kymograph divided by the entire duration of the movie. Only microtubules that showed consistent and measurable growth for the entire duration of the movie were included in the analysis. For determining catastrophe frequencies of γTuRC-nucleated microtubules, the lifetime of each microtubule was also measured (from the point of appearance until it disappears in the kymograph) and the catastrophe frequency of an individual microtubule was calculated as the number of catastrophes divided by the microtubule’s lifetime. The reported catastrophe frequencies are averages of catastrophe frequencies of individual microtubules.

### Fraction of time in growth phase

For “typical” microtubule dynamics, stochastic switching events between growing and shrinking phases were determined manually using a custom MATLAB interface to record the time points of switching by clicking with the computer mouse, beginning with the first instance of observed growth and ending with the last frame (bottom row) of the kymograph or the last frame in which the microtubule end could be distinguished. Only kymographs with at least one recorded switching event were used for quantification, in which the sum of the growth event durations (the difference between the 1^st^ and 3^rd^, 3^rd^ and 5^th^ recorded time points, and so on) was divided by the total measured duration (the difference between the 1^st^ and last timepoints). For lower concentrations of KIF2A (2, 5 nM) and MCAK (5, 10, 20 nM), minus end growth events were difficult to resolve by eye for many kymographs, due to small, fast, and heterogeneous fluctuations closer to the limits of our spatial resolution and frame rate. This precluded using the manual analysis normally performed for the stereotypical, saw-toothed kymographs produced by longer-duration dynamic episodes. Therefore, an automated, custom MATLAB script was used to identify minus ends per-frame, producing a trace of growth. Traces were manually inspected and traces that failed to track the end correctly were discarded before calculating the fraction of growth time. This calculation was performed for each trace by summing the number of frames in which growth was detected over the total number of frames.

### γTuRC-nucleated microtubule analysis

For each nucleation assay, microtubules were counted manually using the ‘Point tool’ in Fiji, at 6 different time points until the end of the movie. A microtubule was considered γTuRC-nucleated if only one end displayed episodes of growth and shrinkage while the other remained capped. At a given time point, the total number of γTuRC-nucleated microtubules was obtained by adding the newly nucleated microtubules to the total number obtained at the previous time point. This quantification included both γTuRC-capped and γTuRC-released microtubules. Each γTuRC-nucleated microtubule that detached from γTuRC at any point of the experiment was counted as γTuRC-released.

Each microtubule that displayed dynamicity at both ends from the first frames of observation was considered a spontaneously-nucleated microtubule. The number of spontaneously nucleated microtubules over time was quantified as described above for the γTuRC-nucleated ones.

A microtubule was considered to be treadmilling when a γTuRC-released microtubule displayed minus end depolymerization and a dynamic plus end, such that the microtubule seemed to translocate along the glass surface.

### γTuRC-mGFP-spastin co-localization analysis

For each co-localization experiment at least five different fields of view were imaged per condition tested. After averaging, fluorescent intensities were measured in the entire fields of view using Fiji.

### Data plots and errors

All data plots were generated using Microsoft Excel or Prism 9 (GraphPad Software, San Diego, CA, USA). Variability of the data is represented by the standard error of the mean (SEM) using vertical error bars. In cases where none of the conditions had visible standard error bars, the standard deviation (SD) was used instead to visibly indicate the degree of data variability. It is stated in the figure legends which type of error is plotted.

