## Supplementary information for "The minus end depolymerase KIF2A drives flux-like treadmilling of γTuRC-uncapped microtubules"

### Supplementary Figures

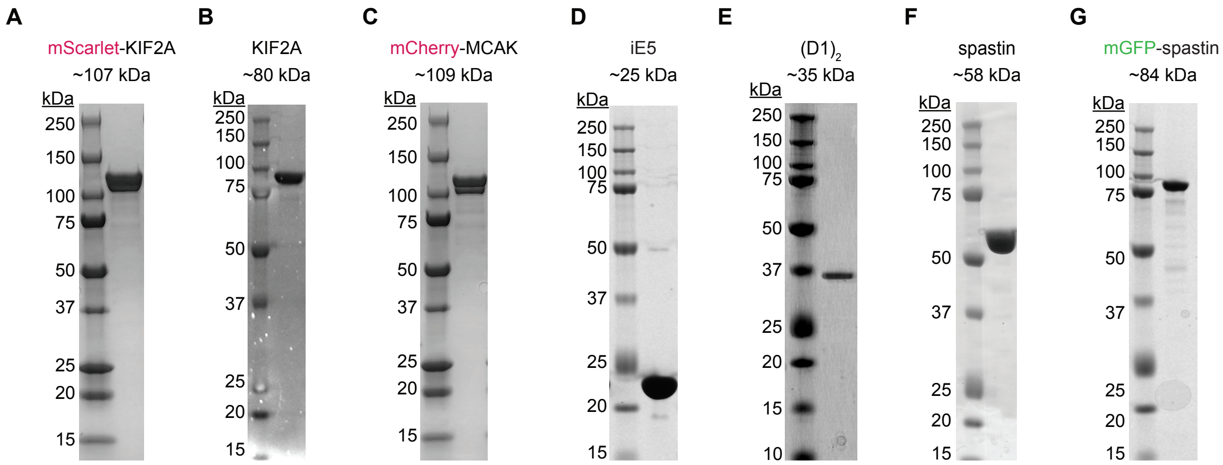


**Supplementary Figure 1. Coomassie-stained SDS gel with purified recombinant proteins used in this study. (A)** mScarlet KIF2A; **(B)** untagged KIF2A; **(C)** mCherry-MCAK; **(D) iE5; (E)** $(D1)_2$; **(F)** spastin; **(G)** mGFP-Spastin. mCherry-tagged proteins migrate as double bands, as previously published (Hentrich and Surrey, 2010; Roostalu et al., 2018), possibly associated with different maturation states of mCherry. mScarlet, derived from mCherry (Bindels et al., 2017), causes a similar migration pattern.

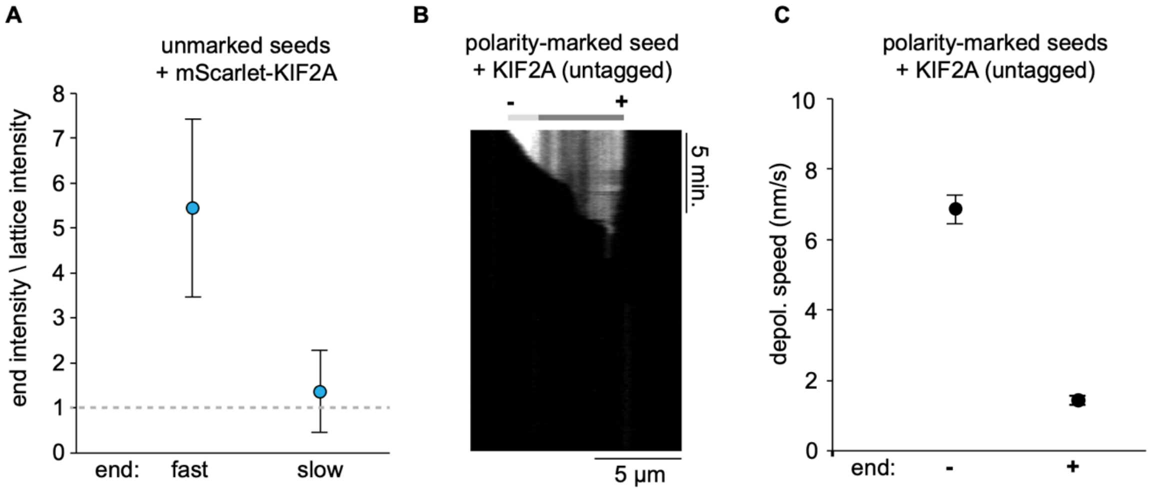


**Supplementary Figure 2. Depolymerization of non-polarity-marked and polarity-marked GMPCPP microtubules by KIF2A. (A)** Ratio of intensities of mScarlet-KIF2A at the fast and slowly growing ends of non-polarity-marked microtubules to its intensity on the microtubule lattice (see Methods). Measurements made from *n* = 27 microtubules. Error bars are SEM. **(B)** Example kymograph demonstrating the minus-end selectivity of KIF2A depolymerzing a polarity-marked GMPCPP microtubule. Polarity as indicated. **(C)** Depolymerization speeds of minus and plus ends of polarity-marked microtubules in the presence of 20 nM untagged KIF2A. Measurements were made from *n* = 69 microtubules. Error bars are SEM.

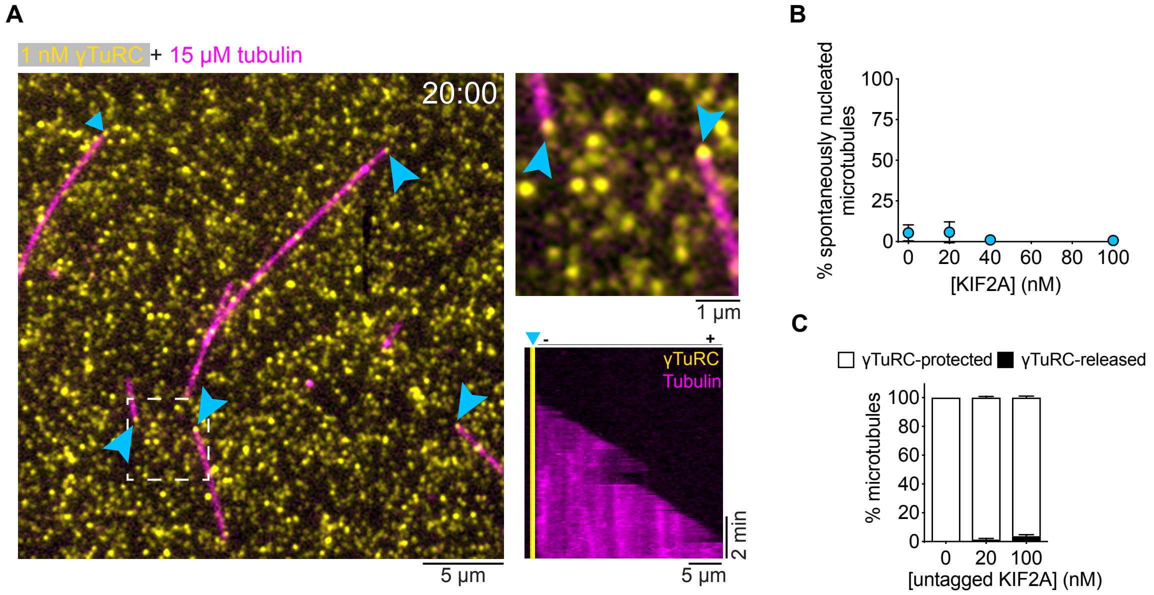


**Supplementary Figure 3. γTuRC-nucleated microtubules in the absence and presence of KIF2A. (A)** (left) TIRF microscopy snapshot of single γTuRC-nucleated microtubules (magenta) 20 minutes after start of imaging, in the presence of 15 µM tubulin (AlexaFluor647, 5,4%). 1 nM biotinylated and BFP-tagged γTuRC (yellow) was used for immobilization. γTuRC signal is an average of all frames imaged during 20 minutes. (right, top) The inset of the dashed area shows two γTuRC-capped microtubules. γTuRC- mediated nucleation sites are indicated by cyan arrows. (right, bottom) Kymograph showing microtubule (magenta) minus-end capping by γTuRC (yellow) and a dynamic plus end. **(B)** Plot of the percentage of spontaneously nucleated microtubules in solution in γTuRC nucleation assays using 12.5 µM of tubulin and different mScarlet-KIF2A (cyan) concentrations, as indicated. Number of microtubules analyzed per condition: n = 256; 20 nM, n = 249; 40 nM, n = 190; 100 nM, n = 91. Data for plots were pooled from three independent experiments. Error bars are SEM. For symbols without visible error bars, error bars are smaller than the symbol size. **(C)** Bar graph of the mean percentage of microtubules that either remain protected by γTuRC or are released after nucleation, in the presence of the indicated concentrations of untagged KIF2A. Number of microtubules analyzed per condition: 0 nM, n = 162; 20 nM, n = 199; 100 nM, n = 83; Data for plots were pooled from at least two independent experiments. Error bars are SEM. 2 nM of γTuRC was used for immobilization in B and C.

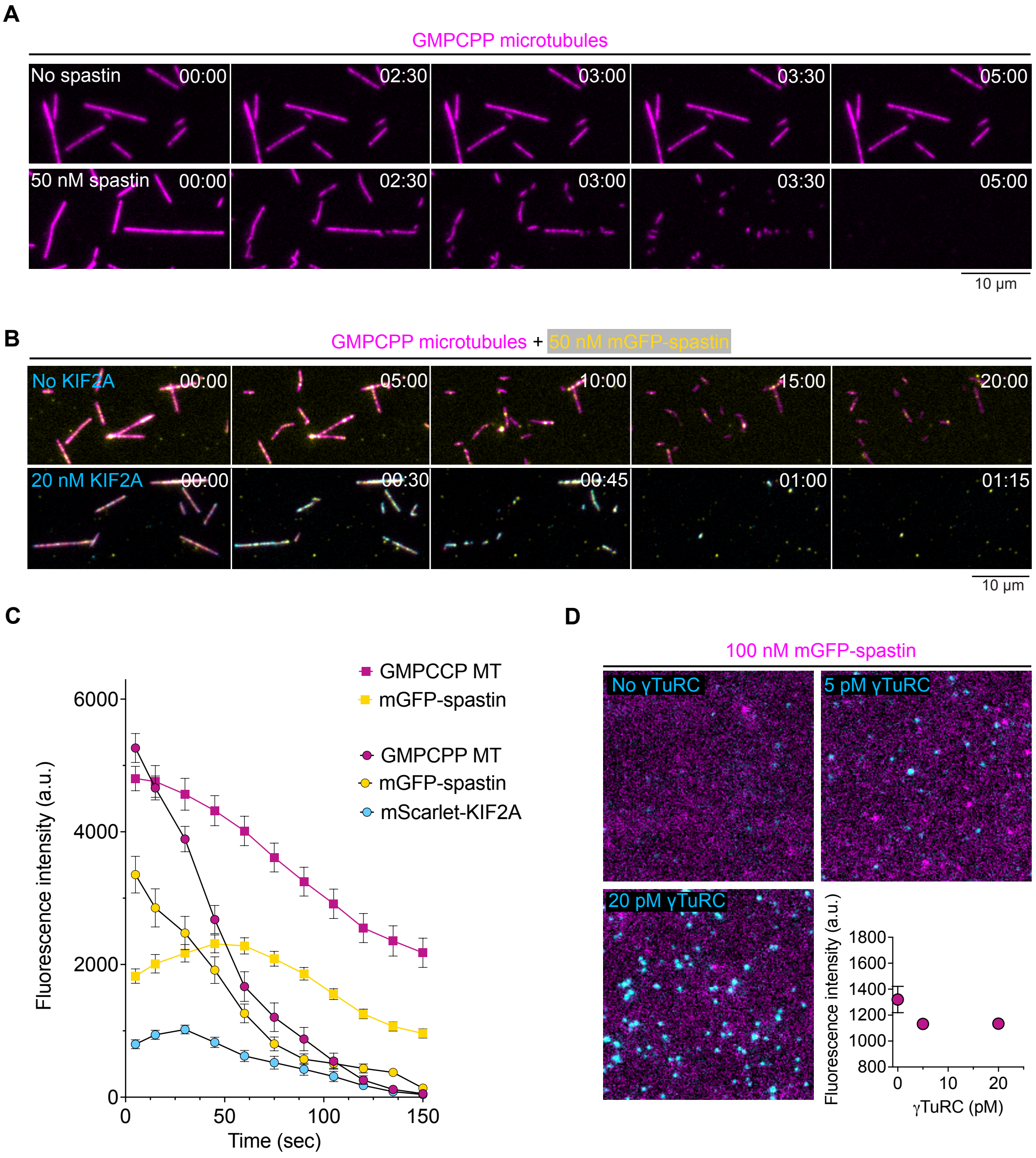


**Supplementary Figure 4. Severing of stabilized microtubule seeds by the severase spastin and mGFP-spastin. (A)** Time sequence of TIRF microscopy images of immobilized GMPCPP microtubules (AlexaFluor647 5%, magenta) in the absence or presence of 50 nM of spastin. **(B)** Time sequence of TIRF microscopy images of immobilized GMPCPP microtubules (Atto647 5%, magenta) and mGFP-spastin (yellow) in the absence or presence of 20 nM of mScarlet-KIF2A (cyan). Time in B and C is min:s. **(C)** Time course of fluorescence intensities of GMPCCP microtubules (magenta), mGFP-spastin (yellow), and mScarlet-KIF2A (cyan)

measured along the initial contour of microtubules in the presence of 50 nM mGFP-spastin (squares) or 50 nM mGFP-spastin and 20 nM KIF2A (circles). Number of microtubules analyzed per condition: 50 nM mGFP-spastin, *n* = 81; 50 nM mGFP-spastin and 20 nM KIF2A, *n* = 87. Data for plots were pooled from at least two independent experiments. Error bars are SEM. For symbols without visible error bars, error bars are smaller than the symbol size. **(D)** TIRF microscopy images of 100 nM mGFP-spastin in the absence or presence of immobilized γTuRC. γTuRC concentrations used for immobilization as indicated. The fluorescence intensities of mGFP-spastin (bottom, right; see Methods) represent the background intensity and are independent of the γTuRC density on the surface, demonstrating absence of interaction between γTuRC and spastin. Moreover, no colocalization is observed, also indicating absence of interaction. Data for plots were pooled from at least two independent experiments. Error bars are SEM. For symbols without visible error bars, error bars are smaller than the symbol size.

**Movie Legends**

**Movie 1. Asymetric depolymerization of stabilized microtubules by KIF2A.** GMPCPP-stabilized microtubules (magenta) undergoing depolymerization in the presence of 20 nM mScarlet-KIF2A (cyan). Scale bar 5 µm. Related to Figure 1.

**Movie 2. Minus end-selective depolymerization of dynamic microtubules by KIF2A.** Dynamic microtubules (magenta) growing from brightly-labeled GMPCPP seeds in the presence of 12.5 µM tubulin, 2 nM (top) and 20 nM (bottom) mScarlet-KIF2A. Scale bars 5 µm. Related to Figure 2.

**Movie 3. KIF2A drives treadmilling of dynamic microtubules.** Treadmilling microtubules growing in the presence of 22.5 µM tubulin from brightly-labeled GMPCPP seeds (magenta) in the presence of 20 nM mScarlet-KIF2A (cyan). Yellow arrows track depolymerizing minus ends of selected treadmilling microtubules. Scale bar 5 µm. Related to Figure 3.

**Movie 4. KIF2A decreases γTuRC-mediated microtubule nucleation.** Microtubules (magenta) nucleated from surface-immobilized γTuRC in the presence of 12.5 µM tubulin and mScarlet-KIF2A (cyan), at the indicated concentrations. Scale bar 10 µm. Movie corresponding to Fig. 4B.

**Movie 5. KIF2A-driven treadmilling event of a γTuRC-nucleated microtubule.** Microtubule (magenta) nucleated from surface-immobilized γTuRC (yellow) in the presence of 12.5 µM tubulin and 20 nM of mScarlet-KIF2A (cyan). After γTuRC- mediated microtubule nucleation, in very rare cases KIF2A disrupts the γTuRC-microtubule interface, triggering microtubule release and subsequent treadmilling. The white arrowhead follows the microtubule minus end. Scale bar 5 μm. Movie related to Fig. 4H and J.

**Movie 6. γTuRC-nucleated microtubules are severed when KIF2A and spastin act in synergy.** Severing of γTuRC-nucleated microtubules (magenta) in the presence of 11 μM tubulin, 100 nM of spastin, and 20 nM of mScarlet-KIF2A (cyan). Scale bar 2 μm. Movie corresponding to Fig. 5B.

**Movie 7. KIF2A and spastin drive the release and treadmilling of γTuRC-nucleated microtubules.** Microtubules (magenta) nucleated from surface-immobilized γTuRC in the presence of 11 μM tubulin, 100 nM spastin and 20 nM of mScarlet-KIF2A (cyan). After γTuRC-mediated microtubule nucleation, microtubule release and subsequent treadmilling is triggered by the combined action of spastin and KIF2A. The yellow arrowheads point to the minus ends of the microtubules. Scale bar 2 μm. Movie corresponding to Fig. 5C.
